# A Conserved Disruption of the Nuclear Permeability Barrier in Meiosis is Controlled by a Kinase-Phosphatase Pair in *Saccharomyces cerevisiae*

**DOI:** 10.1101/2025.05.14.654091

**Authors:** Madison E. Walsh, Keerthana Chetlapalli, Srigokul Upadhyayula, Grant A. King, Elçin Ünal

## Abstract

In eukaryotic organisms, the nucleus is remodeled to accommodate the space required for chromosome segregation. Remodeling strategies range from closed division, where the nuclear envelope remains intact, to open divisions, where the nuclear envelope is temporarily disassembled. While the budding yeast *Saccharomyces cerevisiae* undergoes closed mitosis, its meiotic nuclear division strategy is less understood. Here we investigate nuclear permeability during meiosis in budding yeast and discover that meiosis II represents a semi-closed division marked by bidirectional mixing between the nucleus and cytoplasm. This includes nuclear entry of the Ran GTPase activating protein (RanGAP), typically cytoplasmic, although RanGAP relocalization appears to be a consequence, rather than a cause of permeability changes. This intercompartmental mixing occurs without nuclear envelope breakdown or dispersal of nucleoporins and is independent of known nuclear pore complex remodeling events. This phenomenon, termed virtual nuclear envelope breakdown (vNEBD), represents a unique mechanism distinct from other semi-closed divisions. We demonstrate that vNEBD is integrated into the meiotic program and regulated by the conserved meiotic kinase Ime2 and the meiosis-specific protein phosphatase 1 regulatory subunit, Gip1. Remarkably, the vNEBD event is conserved between *S. cerevisiae* and the distantly related *Schizosaccharomyces pombe*, indicating a conserved, critical role in meiosis.

## Introduction

The integrity of the nucleus and the regulation of nucleocytoplasmic transport are critical for maintaining cellular homeostasis in eukaryotic cells. This compartmentalization between the cytoplasm and nucleoplasm is maintained by the double-membraned nuclear envelope (NE). Embedded within the NE, the nuclear pore complex (NPC) functions as a selective gate, controlling molecular exchange between the nucleus and cytoplasm (reviewed in Lin and Hoelz 2019). While small molecules and ions can passively diffuse through the NPC, macromolecules larger than ∼30 kDa require specialized transport receptors known as karyopherins. NPCs are built from multiple proteins called nucleoporins, which assemble into distinct substructures that confer selectivity to the transport process (reviewed in Lin and Hoelz 2019). Karyopherins mediate the transport of cargo by interacting with channel nucleoporins. Directionality of transport is established by a Ran GTPase gradient – RanGTP in the nucleus and RanGDP in the cytoplasm – regulated by the enzymes RanGEF and RanGAP (Izaurralde et al. 1997; reviewed in Wing, Fung, and Chook 2022). Disruption of nucleocytoplasmic compartmentalization can have severe consequences in humans; for instance, defects at the nuclear periphery have been linked to neurodegenerative disorders such as amyotrophic lateral sclerosis (ALS) and Huntington’s disease (reviewed in Liu and Hetzer 2022; Martins et al. 2020).

An intact nuclear periphery plays a crucial role in maintaining eukaryotic cellular health, yet it undergoes significant remodeling during every cell division to facilitate the accurate segregation of genetic material. To accommodate this process, eukaryotic organisms have evolved different nuclear division strategies (Dey and Baum 2021). At one extreme, complete NE breakdown (NEBD) occurs during cell division (“open” divisions), such as in mitosis in human cells. Most fungi, including yeast, are at the other side of the spectrum, where the NE expands to remain intact throughout the division and nucleocytoplasmic transport is maintained (“closed” division). Notably, many organisms exhibit more intermediate divisions, where part of the NE is maintained but intercompartmental mixing between the nucleoplasm and cytoplasm occurs. These intermediates, termed either “semi-closed” or “semi-open” divisions, typically disrupt nuclear transport via two mechanisms: partial NPC disassembly and/or local NEBD (reviewed in Dey and Baum 2021). For example, the fungus *Aspergillus nidulans* partially disassembles the NPC via phosphorylation (“semi-closed;” De Souza et al. 2004), while the fission yeast *Schizosaccharomyces japonicus* undergoes local NEBD during anaphase in mitosis (“semi-open;” Yam et al. 2011).

Different nuclear division strategies can occur in the same organism depending on cell type or developmental stage, such as during mitosis versus meiosis (reviewed in Walsh, King and Ünal 2024). In the fission yeast *Schizosaccharomyces pombe*, mitosis proceeds via a classical closed division. However, during meiosis II, a striking departure from closed division is observed where mixing between nuclear and cytoplasmic contents occurs (Asakawa et al. 2010; Arai et al. 2010). Notably, electron microscopy reveals no fenestrations in the nuclear envelope (NE), and live-cell imaging of nearly all nucleoporins throughout meiosis shows no signs of NPC disassembly (Asakawa et al. 2010). This unusual form of nucleocytoplasmic exchange has been termed “virtual nuclear envelope breakdown” (vNEBD) to reflect its distinct mechanism (Flor-Parra et al. 2018). The budding yeast *Saccharomyces cerevisiae*, which diverged from *S. pombe* over 400 million years ago (Sipiczki 2000), also undergoes closed mitosis. Intriguingly, a recent study in *S. cerevisiae* observed the relocalization of two nuclear proteins to the cytoplasm specifically during meiosis II (Shelton et al. 2021). Although the molecular basis of this relocalization remains poorly understood, it raises the possibility that *S. cerevisiae* might also experience a form of global nucleocytoplasmic mixing during meiosis—potentially analogous to vNEBD.

A growing body of research has demonstrated that the nucleus undergoes significant remodeling during *S. cerevisiae* meiosis. During meiosis I, the NPCs exhibit transient, phosphorylation-driven detachment of the nuclear baskets (King et al. 2022). During meiosis II, the nucleus undergoes a five-way division, resulting in the formation of four gamete nuclei and a fifth nuclear compartment, termed the “<u>G</u>ametogenesis <u>U</u>ninherited <u>N</u>uclear <u>C</u>ompartment” (GUNC). This process facilitates the mass turnover of NPCs and contributes to the elimination of age-induced damage (King et al. 2019). Concurrently, mitochondria detach from the cellular periphery, forming extensive membrane contacts with the nucleus (Sawyer et al. 2019). These events are tightly coordinated within the meiotic program and are often regulated by core cell cycle machinery. Despite extensive remodeling, the nuclear envelope remains intact (Moens 1971; Peter B. Moens and Rapport 1971; King et al. 2019) and the channel nucleoporins do not dissociate from the NPC core (King et al. 2019), leaving it unclear whether and how nuclear transport is altered during budding yeast meiosis.

In this study, we investigate nuclear permeability during meiosis in *S. cerevisiae*. We find that, as in *S. pombe*, the nuclear permeability barrier is transiently disrupted in meiosis II, in a large-scale mixing event. This vNEBD event does not require the relocalization of RanGAP or meiotic NPC remodeling. Instead, it is tightly coupled to meiotic progression through key cell cycle regulators: initiation of vNEBD requires the cyclin-dependent kinase (CDK)-like kinase Ime2, whereas its completion depends on the Protein phosphatase 1 (PP1/Glc7) regulatory subunit Gip1. Our findings establish the disruption of nuclear transport as a conserved feature of meiosis, with potentially profound implications for gamete health.

## Results

### *S. cerevisiae* undergoes vNEBD during meiosis II

To investigate whether *S. cerevisiae* undergoes a global mixing event between nucleus and cytoplasm during meiosis, we used live cell imaging to monitor the localization of three different endogenous nucleoplasmic proteins: Npl3, an RNA binding protein (Windgassen et al. 2004; Dermody et al. 2008; Kress, Krogan, and Guthrie 2008; Moehle et al. 2012; Pérez-Martínez et al. 2020); Trz1, a 3’ RNA processing protein (Chen et al. 2005); and Pus1, a pseudouridine synthase (Motorin et al. 1998). All three proteins remained prominently nuclear during meiotic prophase and throughout meiosis I. However, each protein dispersed from the nucleus in late anaphase II and accumulated back in the nucleus shortly after (Fig. 1A, S1A-B). To confirm the generalizability of our results, we examined a fully synthetic reporter containing two copies of the fluorophore mCherry fused with the SV40 nuclear localization signal (NLS). This reporter also exhibited transient dispersal from the nucleus during meiosis II (Fig. 1B). These data are consistent with a previous study reporting Lys21-GFP and Pus1-GFP losing nuclear localization at a similar stage in meiosis (Shelton et al. 2021). Notably, the proteins that we examined span a range of molecular weights (57-125 kDa) and utilize distinct karyopherin import pathways (Srp1/Kap95 for SV40NLS and Mtr10 for Npl3 (Conti et al. 1998; Truant and Cullen 1999; Senger et al. 1998).

**Figure 1.**
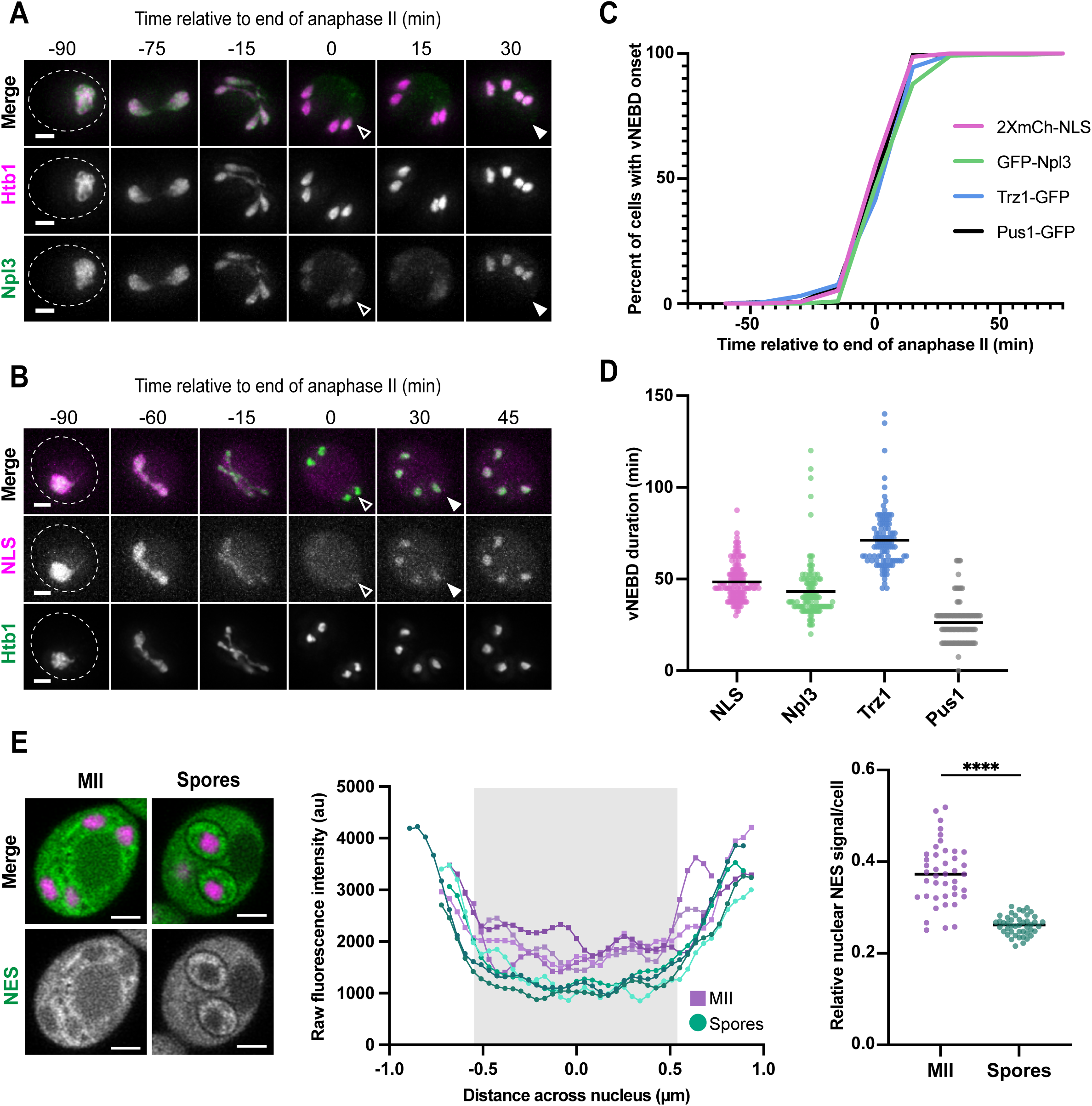
Budding yeast undergoes vNEBD during meiosis II. A-B. Time-lapse microscopy of a cell progressing through meiosis. Open arrows indicate the start of vNEBD, and closed arrows represent the end of vNEBD. Maximum intensity projection shown. **A.** Cells contain fluorescently tagged histone Htb1-mCherry (pink) and nucleoplasmic protein GFP-Npl3 (green) (strain ÜB18509). **B.** Cells contain nucleoplasmic reporter 2XmCherry-SV40NLS (NLS, pink) and fluorescently-tagged histone Htb1-eGFP (green) (strain ÜB21380). **C.** The percent of cells that have initiated vNEBD (proteins disperse from the nucleus) over time (min) relative to the end of anaphase II (first time point of maximum chromatin separation). 2XmCherry-SV40NLS (pink), GFP-Npl3 (green), Trz1-GFP (blue), and Pus1-GFP (gray). Two replicates per strain, 65 ≤ n ≤ 140 cells per replicate. **D.** Duration of vNEBD (nuclear protein dispersal) measured via live time-lapse microscopy (two replicates per strain, 65 ≤ n ≤ 140 cells per replicate). Line represents mean duration (2XmCherry-SV40NLS = 45.5 min, GFP- Npl3 = 43.2 min, Trz1-GFP = 71.2 min, Pus1-GFP = 26.3 min). **E.** (Left) Live Airyscan microscopy of cells during meiosis II (MII) or during post-meiotic spore development expressing histone marker Htb1-mCherry (pink) and PKI-NES-3XeGFP (NES, green) (strain ÜB38411). Single Z-slice shown. (Middle) Line scans from the cells depicted to the left of raw fluorescence intensities (arbitrary units). Purple lines represent scans from each nucleus in the MII cell and teal lines represent scans from each nucleus in the cell with spores. Distance across the nucleus is measured in µm with the gray box representing the approximate nuclear boundary. (Right) Quantification of nuclear NES- 3XeGFP signal in cells staged at late MII (n = 42 cells) or immature (pre-ascal collapse) spores (n = 45 cells). Line scans measured signal intensity across each nucleus, and the ratio of minimum to maximum intensity per nucleus was averaged for each cell. Mann-Whitney test, p<0.001. Scale bar is 2 µm for all images shown. Full sample size information can be found in the Materials and Methods.

The onset of nuclear signal dispersal was highly consistent among the different proteins tested, occurring near the end of anaphase II (Fig. 1C; anaphase II occurring on average 4.8 minutes before 2XmCherry-SV40NLS dispersal, 7.6 minutes before GFP-Npl3 dispersal, 5.1 minutes before Trz1-GFP dispersal, and 6.6 minutes before Pus1-GFP dispersal) and lasting around 45 minutes (Fig. 1D; average durations of 48.5 minutes for 2XmCherry-SV40NLS, 43.2 minutes for GFP-Npl3, 71.2 minutes for Trz1- GFP, and 26.3 minutes for Pus1-GFP). The variation in duration did not correlate with molecular weight of reporters (Fig. S1C). Importantly, quantification of whole-cell fluorescence did not reveal a decrease in fluorescence intensity during the time of nuclear signal dispersal, suggesting that the proteins are relocalizing rather than undergoing mass degradation and resynthesis (Fig. S1D). These data indicate that nuclear proteins undergo a global and transient relocalization event during a discrete time window in meiosis.

We next assessed whether the apparent disruption of nucleocytoplasmic transport occurs in a bidirectional manner by examining the localization of a preferentially cytoplasmic protein. We generated a synthetic reporter consisting of three copies of eGFP tagged with a protein kinase A inhibitor nuclear export signal (PKI-NES- 3xEGFP; Shaikhqasem et al. 2020) and assessed its localization using high resolution Airyscan microscopy in live cells. The nuclear PKI-NES-3xEGFP signal was significantly higher in meiosis II relative to post-meiotic spores (Fig. 1E; Mann-Whiteney test, p < 0.0001), consistent with the cytoplasmic reporter leaking into the nucleus. The increase in nuclear PKI-NES-3xEGFP signal also coincided with nucleoplasmic protein dispersal (Fig. S1E), suggesting the existence of a single, bi-directional mixing event.

Despite these large-scale changes to protein localization, the NE appears to remain largely intact as determined by both serial transmission electron microscopy (P. B. Moens 1971; Peter B. Moens and Rapport 1971; King et al. 2019) and fluorescent microscopy (Fig. S1F). Consistently, we also observed that there were limits to the intermixing between nucleoplasm and cytoplasm. Notably, a dim nuclear signal was still detectable even after the dispersal of NLS-containing proteins (Fig. 1A, S1B). Likewise, the PKI-NES-3xEGFP reporter, despite exhibiting some entry into the nucleus, was still enriched in the cytoplasm (Fig. 1E, left). Furthermore, ribosomal proteins remained robustly excluded from the nucleus throughout meiosis II (Fig. S1G). Intermixing of the cytoplasm and nucleoplasm, therefore, does not appear to be facilitated by complete NEBD or sizable fenestrations in the NE that would result in uniform mixing and allow for large structures like the ribosomes to freely enter the nucleus. We therefore established that *S. cerevisiae* undergoes a vNEBD event during meiosis II, similar to that observed in the distantly related fission yeast *S. pombe*.

### RanGAP relocalization does not drive vNEBD

Because the NE appears to remain largely intact throughout meiosis, we next investigated whether vNEBD is driven by changes in the directional transport machinery. Directional transport into or out of the nucleus is achieved by the asymmetric distribution of Ran-GTP and Ran-GDP, which facilitate nuclear import and export respectively. This gradient is maintained by the nuclear-localized Ran Guanine Exchange Factor (RanGEF; called Prp20 in budding yeast), which stimulates the release of GDP from Ran to allow binding of GTP, and the cytoplasmic Ran GTPase activating protein (RanGAP; Rna1 in budding yeast), which accelerates the hydrolysis of Ran-GTP to Ran-GDP (Fig. 2A; reviewed in Wing, Fung, and Chook 2022). We first sought to determine whether these key transport regulators exhibit altered localization during vNEBD. We monitored the localization of Prp20 and Rna1 in live meiotic cells. The RanGEF Prp20 remained co-localized with histone Htb1 throughout both meiotic divisions (Fig. 2B), consistent with its known chromatin association (Clark et al. 1991; Akhtar et al. 2001). In contrast, the RanGAP Rna1 appeared to enter the nucleus during late meiosis II, despite its strong exclusion from the nucleus during meiosis I and post meiosis II (Fig. 2C, S2A). This relocalization could disrupt the asymmetric distribution of Ran-GTP and Ran-GDP, potentially driving vNEBD.

**Figure 2.**
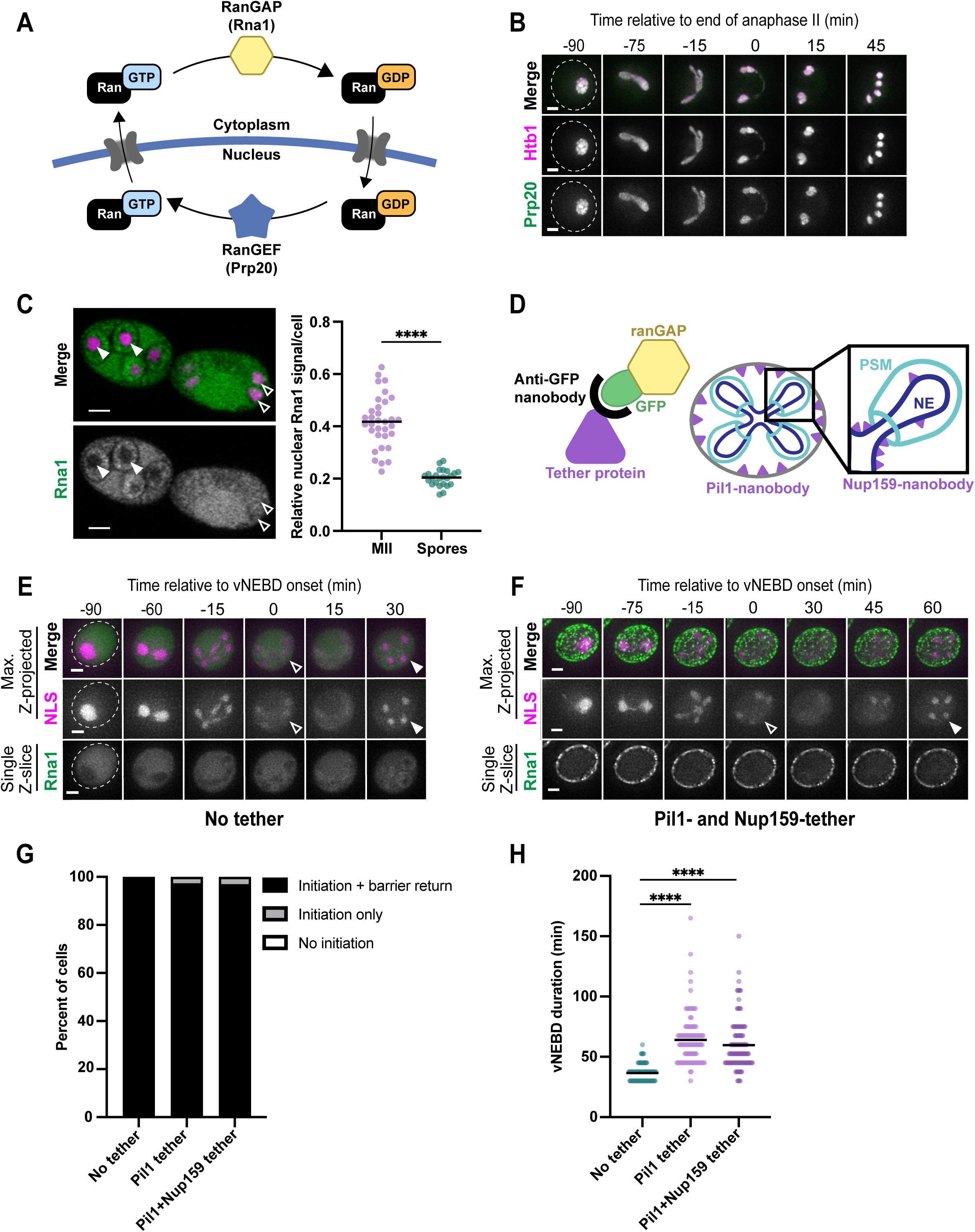
RanGAP relocalization to the nucleus does not drive vNEBD. **A.** Schematic representing how RanGEF and RanGAP establish asymmetric populations of RanGTP and RanGDP essential for directional nucleocytoplasmic transport. NPCs are represented in gray. **B.** Time-lapse microscopy of a cell progressing through meiosis tagged with histone marker Htb1-mCherry (pink) and RanGEF Prp20-GFP (green) (strain ÜB20161). Maximum intensity projection shown. **C.** (Left) Live Airyscan microscopy of sporulating cells in MII (right cell) and immature spores (left cell) expressing histone marker Htb1-mCherry (pink) and RanGAP Rna1-3XeGFP (green) (strain ÜB20153). Single Z-slice shown. Arrowheads point to nuclei in-focus, with closed arrows indicating exclusion of Rna1 from the nucleus and open arrows indicating nuclear Rna1 signal. (Right) Quantification of nuclear Rna1 signal in cells staged at late MII (n = 33 cells) or immature (pre-ascal collapse) spores (n = 24 cells). Line scans measured signal intensity across each nucleus, and an average of the ratio of minimum to maximum intensity was taken for each cell, such that each point represents one cell. Mann-Whitney test, p<0.001. **D.** Schematic representing the experimental strategy to tether Rna1 outside of the nucleus. (Left) Tether protein (purple) is fused to an anti-GFP nanobody (black), which can bind to the 3XeGFP tag (green) fused to Rna1 (yellow). (Right) Schematic of the dividing nucleus with the NE in dark blue and developing gamete plasma membranes (teal) typical of late anaphase II. The Pil1 tether (purple) localizes to the cell periphery, outside of the developing gametes, whereas Nup159 is localized to the outside of the NE. **E-F.** Time-lapse microscopy of a cell progressing through meiosis tagged with histone marker Htb1-mCherry (pink) and RanGAP Rna1- 3XeGFP (green) with either no tether protein (strain ÜB20155) (**E**), or both Pil1 and Nup159 tether proteins (strain ÜB36287) (**F**). Open arrows indicate the start of vNEBD, and closed arrows represent the end of vNEBD. The first two rows show a maximum intensity projection, and the last row shows a single z-slice. **G**. The percentage of cells that underwent meiosis II (MII) with vNEBD initiation and barrier return (black), vNEBD initiation with no reestablishment of the permeability barrier (gray), or no vNEBD initiation (white, no cells). Pil1+Nup159 tether refers to cells containing both the Pil1- nanobody and the Nup159-nanobody. Two replicates scored for each category (79 ≤ n ≤ 120 cells per replicate). **H.** Duration of vNEBD in minutes as assessed by 15-minute interval time-lapse microscopy. Each point represents one cell. Black line represents the mean vNEBD duration (no tether = 36.5 min, Pil1-tether = 64.1 min, Pil1- and Nup159- tether = 59.7 min). For each condition, two replicates were scored (79 ≤ n ≤ 120 cells per replicate). Scale bar is 2 µm for all images shown. Full sample size information can be found in the Materials and Methods.

RanGAP similarly enters the nucleus during vNEBD in *S. pombe* (Arai et al. 2010; Asakawa et al. 2010) and its forced localization into the nucleus produces a similar phenotype to vNEBD, indicating that nuclear RanGAP is sufficient to disrupt nuclear permeability barrier (Asakawa et al. 2010). However, it remains unclear whether RanGAP’s entry into the nucleus initiates vNEBD or whether it is instead a consequence of vNEBD. To distinguish between these two possibilities, we asked whether vNEBD could still occur if Rna1 was constitutively kept outside of the nucleus throughout meiosis (Fig. 2D). We forced cytoplasmic localization of Rna1-3xGFP by using anti-GFP nanobodies fused to either Pil1, an eisosomal protein that localizes at the plasma membrane (Walther et al. 2006), or to both Pil and the cytoplasmic-facing nucleoporin Nup159 (Hinshaw 1994). We observed robust cytoplasmic retention of RanGAP under these conditions (Fig. 2F). However, vNEBD still occurred with similar penetrance and timing relative to control samples (Fig. 2E-G). Therefore, RanGAP’s entry into the nucleus is not necessary for vNEBD and instead represents a consequence of general nucleocytoplasmic mixing.

While tethering Rna1 constitutively outside of the nucleus did not influence vNEBD onset, it did affect the duration of vNEBD. vNEBD duration was significantly longer when Rna1 was tethered to either Pil1 alone or Pil1 and Nup159 relative to the control (Fig. 2H; 36.5 minutes for untethered cells, compared to 64 minutes for Pil1- tethered cells and 59.7 minutes for Pil1- and Nup159-tethered cells; Mann-Whiteney test, p < 0.001). Rna1 tethered to Pil1 is trapped outside of the developing spores after prospore membrane closure (Fig. 2D, F). This reduced access to Rna1 may delay re-establishment of the Ran-gradient and selective nucleocytoplasmic transport. Further, spore nuclei from the same progenitor cell exhibited greater heterogeneity in vNEBD duration when Rna1-GFP was tethered (Fig. S2B-D), consistent with the possibility that different sized pools of Rna1 are being inherited. However, these spores were still able to grow and divide when supplied with nutrients (Fig. S2E), indicating that lengthy vNEBD does not noticeably compromise spore health.

### vNEBD is precisely timed relative to other meiotic nuclear remodeling events

To understand how vNEBD might be linked to or impacted by different meiotic events, we examined the correlation between the vNEBD start and finish with characteristic nuclear events that occur during meiosis (Fig. 3A), including: NPC sequestration (Nup49; Fig. 3B) and nuclear basket return (Nup60; Fig. 3C); mitochondrial collapse onto the nuclear envelope (Cit1; Fig. S3A); nucleolar sequestration and release of a cell-cycle phosphatase (Cdc14; Visintin, Hwang, and Amon 1999; Fig. S3B); spindle assembly and disassembly (Tub1; Fig. 3D); onset and completion of chromosome segregation (visualized with Htb1; Fig. 1A, B); and prospore membrane closure (Spo20; Diamond et al. 2009; Fig. 3E). This benchmarking provides an important resource for mechanistic characterization of nuclear remodeling events in budding yeast. vNEBD took place with reproducible timing relative to the other meiotic nuclear remodeling events that were examined, consistent with it representing a tightly regulated part of the meiotic program (Fig. 3F).

**Figure 3.**
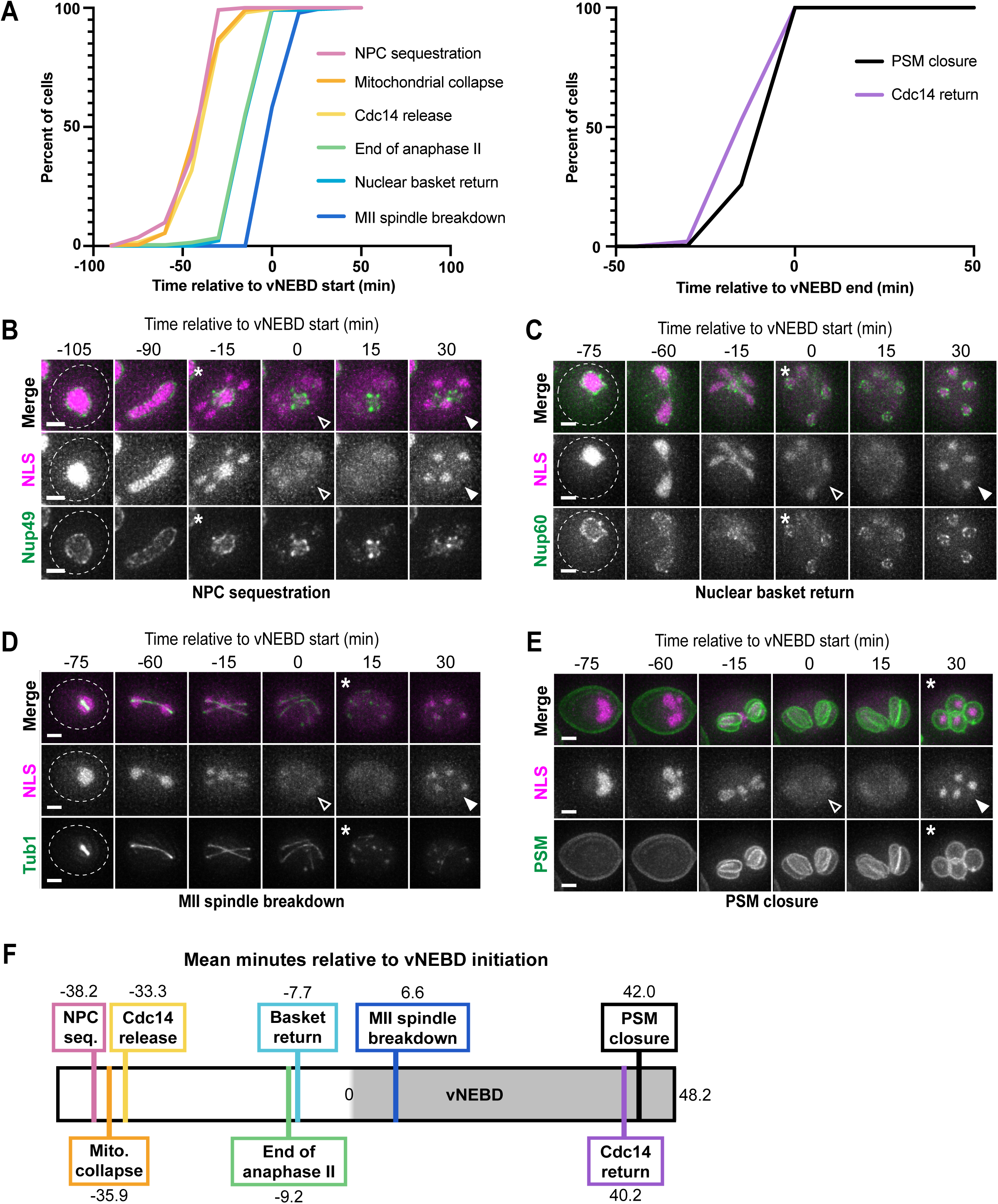
vNEBD is precisely timed relative to other meiotic events. **A**. Timing of key meiotic events relative to vNEBD initiation (left) and conclusion (right). Events timed around vNEBD onset are NPC sequestration to the GUNC (pink), mitochondrial collapse onto the nucleus (orange), Cdc14 release from the nucleolus (yellow), the end of anaphase II (green), nuclear basket return (light blue), and meiosis II spindle breakdown (dark blue). Two replicates each strain with 81≤ n ≤ 154 cells per replicate. Events timed around vNEBD end are the closure of the prospore membranes (black) and return of Cdc14 to the nucleolus (purple). Two replicates are measured for each strain with 111 ≤ n ≤ 137 cells per replicate. **B-E.** Time-lapse microscopy of a cell progressing through meiosis. Open arrows indicate the start of vNEBD, and closed arrows represent the end of vNEBD. Maximum intensity projection shown. Scale bar is 2 µm. **B.** Cells contain reporter 2XmCherry-SV40NLS (pink) and fluorescently tagged channel FG-nucleoporin Nup49-GFP (green) (strain ÜB18513). Asterisk marks the onset of NPC sequestration away from the forming gamete nuclei. **C.** Cells contain reporter 2XmCherry-SV40NLS (pink) and GFP-tagged nuclear basket protein Nup60- GFP (green) (strain ÜB44905). Asterisk marks the first time point of nuclear basket return to the gamete nuclei. **D.** Cells contain reporter 2XmCherry-SV40NLS and exogenous fluorescently tagged tubulin Tub1-GFP (strain ÜB33507). Asterisk marks first time of meiosis II spindle breakdown. **E.** Cells contain reporter 2XmCherry- SV40NLS (pink) and prospore membrane marker (PSM; spo20(51-91)-eGFP) (strain ÜB34583). Asterisk marks the first time point of PSM closure. **F.** Schematic showing the relative order of meiotic events to-scale using the time difference of each relative to vNEBD onset or end. Numbers represent the mean time difference from vNEBD onset (t = 0) in minutes. Full sample size information can be found in the Materials and Methods.

Of note, vNEBD onset was coincident with meiosis II spindle disassembly (Fig. 3A, D). In fact, we never observed spindle disassembly prior to vNEBD start (n = 235 cells). In *S. pombe*, vNEBD has been proposed to play a role in timely spindle breakdown in meiosis II by bypassing the need for active import of disassembly factors into the nucleus (Flor-Parra et al. 2018). Cells deficient in vNEBD through deletion of fission-yeast specific gene *SPO5* develop hyper-extended spindles in meiosis II, but spindle length is unaffected in meiosis I (Asakawa et al. 2010; Flor-Parra et al. 2018). The temporal correlation of vNEBD and meiosis II spindle breakdown in *S. cerevisiae* may indicate that spindle breakdown timing could be facilitated by vNEBD in meiosis II as it is in *S. pombe*, though further work is necessary to establish a causative relationship.

### vNEBD can be uncoupled from known NPC remodeling events

NPC remodeling underlies changes in nuclear permeability in other organisms that undergo a semi-closed division in which the NE remains intact (reviewed in Walsh, King, and Ünal 2024). For example, in the fungus *Aspergillus nidulans*, phosphorylation by NIMA kinase drives the dissociation of channel nucleoporins from the core NPC structure, turning the NPCs into tunnels rather than gates, which allows the free exchange of cytoplasm and nucleoplasm (De Souza et al. 2004). However, unlike *A. nidulans*, there is no evidence of channel nucleoporins dissociating from the core NPC structure during meiosis (King et al. 2019). Instead, *S. cerevisiae* undergoes two major NPC remodeling events in meiosis II with timing similar to key vNEBD events: NPC sequestration, which occurs on average 38 minutes prior to vNEBD onset (Fig. 3B), and nuclear basket return, which coincides with vNEBD onset (Fig. 3C). In brief, core nucleoporins are sequestered to the GUNC outside the four gamete nuclei during anaphase II, a compartment that is subsequently destroyed during spore maturation (Fig. 4A, left; King et al. 2019). The entire nuclear basket detaches from the core NPC and returns to the gamete nuclear periphery (Fig. 4A, right).

**Figure 4.**
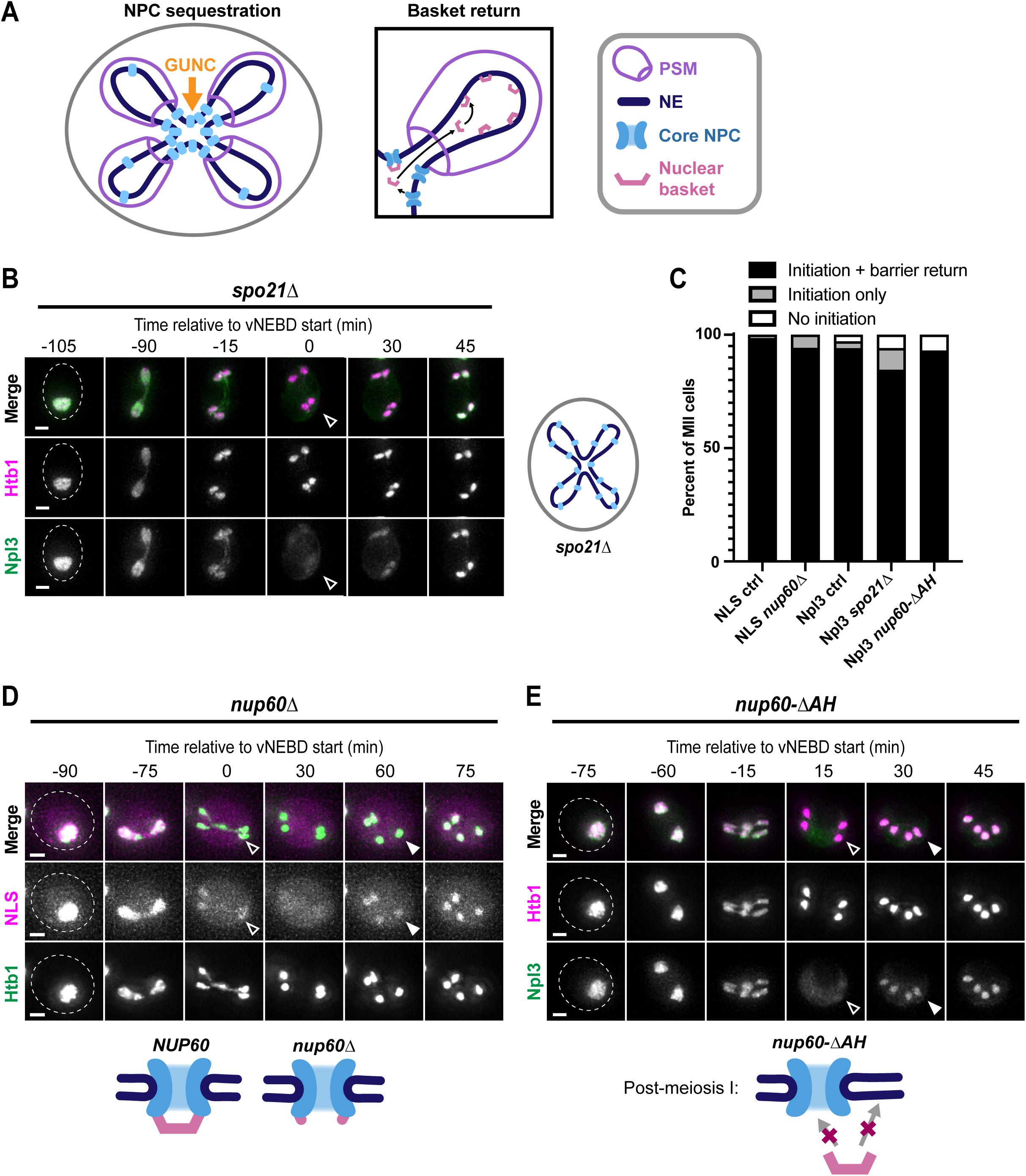
vNEBD can be uncoupled from known NPC remodeling events. **A.** Schematic diagramming the two NPC remodeling events in meiosis II. (Left) Most NPCs (blue) are excluded from growing prospore membranes (PSM; purple), resulting in their sequestration to the GUNC (orange arrow). (Right) Following sequestration to the GUNC, the nuclear basket (pink) detaches from the NPC core and returns to the gamete nuclei, where they associate with the nuclear periphery. **B, D-E.** Time-lapse microscopy of a cell progressing through meiosis. Open arrows indicate the start of vNEBD, and closed arrows represent the end of vNEBD. Maximum intensity projection shown. Scale bar 2 µm. **B.** (Left) Cells contain fluorescently tagged histone Htb1- mCherry (pink) and nucleoplasmic protein GFP-Npl3 in the mutant *spo21*Δ background (strain ÜB21612). (Right) Cartoon showing that NPCs fail to be sequestered to the GUNC when PSMs do not form in *spo21*Δ mutants. **C.** The percentage of cells that underwent meiosis II (MII) with vNEBD initiation and barrier return (black), vNEBD initiation with no reestablishment of the permeability barrier (gray), or no vNEBD initiation (white) for various mutant backgrounds. NLS ctrl (strain ÜB21380) and *nup60*Δ (strain ÜB25072) both contain Htb1-eGFP and 2XmCherry-SV40NLS. *GFP-NPL3* ctrl (strain ÜB18509), *spo21*Δ (strain ÜB36318), and *nup60*Δ*AH* (strain ÜB25839) all contain Htb1-mCherry and GFP-Npl3. Two replicates used for each strain (62 ≤ n ≤ 118 cells per replicate). **D.** (Top) A cell containing reporter 2XmCherry-SV40NLS (pink) and fluorescently tagged Htb1-eGFP (green) in the mutant *nup60*Δ background (strain ÜB25072). (Bottom) Cartoon representing the state of the nuclear basket in wild-type *NUP60* cells (left) with an intact basket or *nup60*Δ cells (right), which cannot assemble four of the five basket nucleoporins. **E.** (Top) A cell containing fluorescently tagged histone Htb1-mCherry (pink) and nucleoplasmic protein GFP-Npl3 (green) in the mutant *nup60-*Δ*AH* background (strain ÜB25839). (Bottom) Cartoon representing the state of the nuclear basket in nup60-ΔAH cells, in which the nuclear basket cannot return to the nuclear periphery following meiosis. Full sample size information can be found in the Materials and Methods.

We sought to determine whether vNEBD is mechanistically coupled to either of the two meiosis II NPC remodeling events. First, to test whether vNEBD onset is linked to NPC sequestration, we leveraged the fact that NPC sequestration to the GUNC requires formation of prospore membranes (PSMs; King et al. 2019). By disrupting prospore membrane formation (*spo21*Δ), we can force NPCs to remain around developing gamete nuclei. We monitored whether vNEBD was impacted in the absence of NPC sequestration. Surprisingly, vNEBD still took place in *spo21*Δ cells (Fig. 4B-C), with the onset of vNEBD taking place with wild-type timing (Fig. S4B). *spo21*Δ cells did exhibit a mild increase in vNEBD duration (Fig. S4A), although reestablishment of nuclear permeability barrier still took place in almost all cells. vNEBD initiation, therefore, occurs independently of NPC sequestration and PSM formation.

To test whether vNEBD is linked to nuclear basket return to nascent spore nuclei, we utilized mutants of the key nuclear basket nucleoporin Nup60. Nup60 is required for the assembly of three basket nucleoporins (Mlp1, Mlp2, and Nup2) at the NPC (Cibulka et al. 2022; Dilworth et al. 2001; Feuerbach et al. 2002; Palancade et al. 2005) and stabilizes association of the fourth (Nup1; King et al. 2022). Further, Nup60 is necessary for the reassociation of the nuclear basket with the nuclear periphery following meiosis-driven detachment (King et al. 2022). Deletion of an amphipathic helix that interacts directly with the NE perturbs reassociation of the entire nuclear basket with the nuclear periphery in gamete nuclei. We therefore tested whether vNEBD was disrupted by either deletion of the entire *NUP60* gene (*nup60*Δ, complete nuclear basket disruption) or just its amphipathic helix (*nup60-*Δ*AH,* disruption of meiotic nuclear basket return). In both nuclear basket mutants, we observed that almost all cells that underwent meiosis II also experienced vNEBD (Fig. 4C-E; 98.5% for *NUP60*, 94.2% for *nup60*Δ, 93.0% for *nup60-*Δ*AH*) with normal timing relative to the end anaphase II (Fig. S4B). We did observe prolonged vNEBD in *nup60*Δ cells (Fig. S4A), although this may be an indirect effect of low sporulation efficiency and delayed meiotic progression that occurs in *nup60*Δ mutants (Komachi and Burgess 2022). Together, these data suggest that vNEBD can be uncoupled from both major NPC remodeling events in meiosis II, suggesting that vNEBD represents a distinct meiotic event.

### vNEBD initiation is regulated by Ndt80 and Ime2 as part of the meiotic program

Given that the timing of vNEBD is tightly coupled to the meiotic program, we reasoned that it might be controlled by key cell cycle regulators. We first tested whether vNEBD required Ndt80, a core transcription factor that is required for initiating the meiotic divisions (Xu et al. 1995; Hepworth, Friesen, and Segall 1998). We found that vNEBD does not occur in *ndt80*Δ cells, indicating that the Ndt80 transcriptional program is critical for vNEBD (Fig. 5A, C). Since Ndt80 induces the expression of multiple B-type cyclins necessary for the activation of Cdc28, the main cyclin-dependent kinase (CDK) in budding yeast (Reed and Wittenberg 1990), we asked whether Cdc28 activity is required for vNEBD. To specifically test for a role of Cdc28 in meiosis II, we used a conditional allele (*cdc28-as*) that can be selectively inhibited by the ATP analog 1-NM- PP1 (Bishop et al. 2000). Blocking Cdc28 activity prior to meiosis II successfully prevented gamete development; however, cells were still able to progress through other parts of the meiotic program, such as mitochondrial detachment from the cell cortex (Fig. S5A-B). Unexpectedly, blocking Cdc28 activity prior to meiosis II did not disrupt vNEBD initiation and only partially affected vNEBD completion (Fig. 5B-C). The incomplete penetrance of this effect indicates that CDK activity is not strictly necessary for vNEBD completion, may rather play an indirect role, or is involved in a redundant pathway. We also found that cells lacking the meiosis II-specific cyclin Clb3 (*clb3*Δ) exhibited normal vNEBD (Fig. S5C; reviewed in MacKenzie and Lacefield 2020).

**Figure 5.**
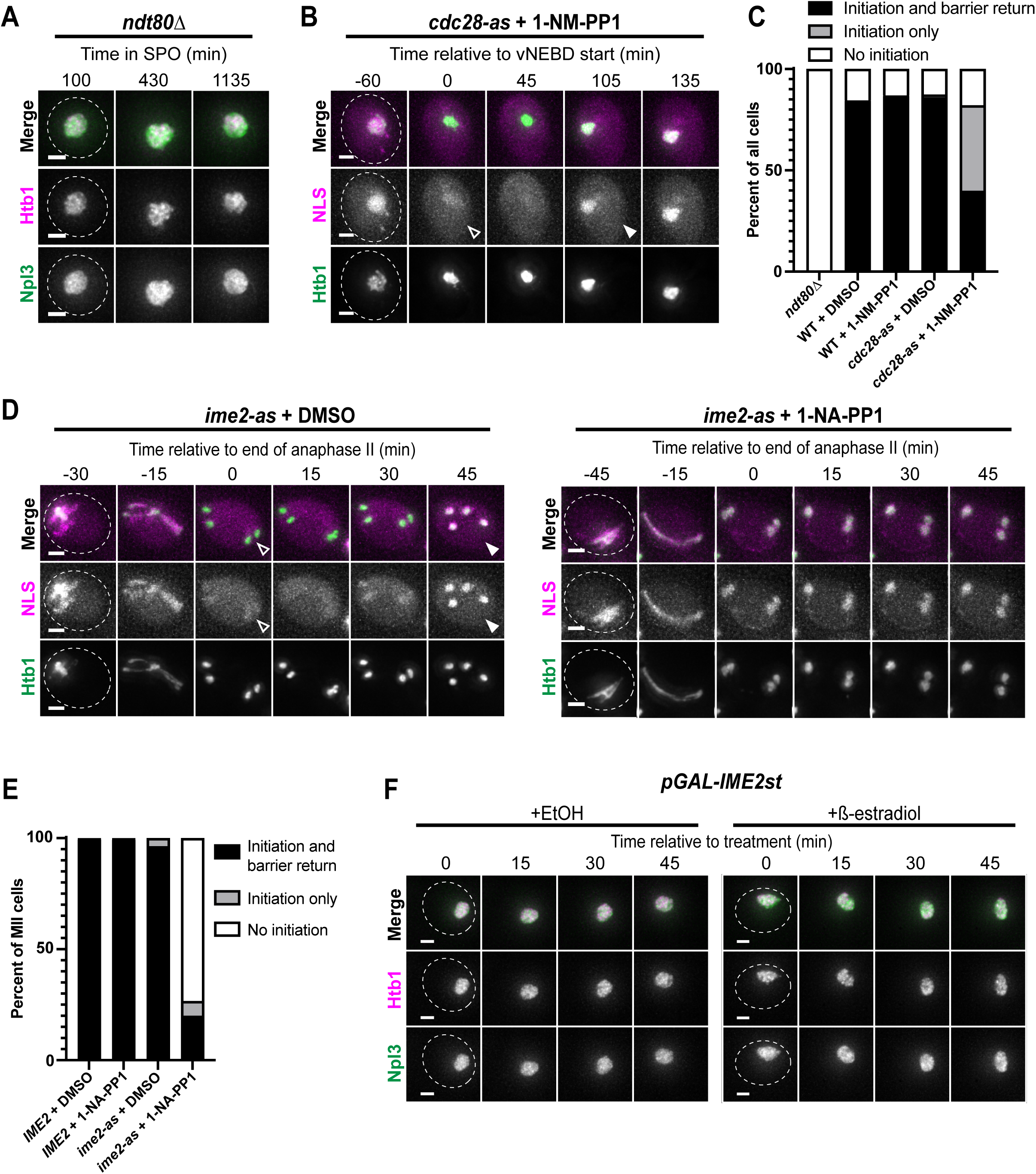
vNEBD initiation is regulated by meiotic cell cycle regulators Ndt80 and Ime2. **A.** Time-lapse microscopy of an arrested *ndt80*Δ cell. Cell contains fluorescently tagged histone Htb1-mCherry (pink) and nucleoplasmic protein GFP-Npl3 (green) (strain ÜB27968). The number of hours in sporulation media is indicated in minutes. **B.** Time-lapse microscopy of a *cdc28-as* cell treated with 1-NM-PP1 as in B. Cell contains reporter 2XmCherry-SV40NLS (pink) and fluorescently tagged histone Htb1-eGFP (green) (strain ÜB36314). **C**. Percentage of all cells in sporulation media with vNEBD initiation and barrier return (black), vNEBD initiation only (gray), or no vNEBD initiation (white). Either DMSO (vehicle) or chemical inhibitor 1-NM-PP1 was added to a final concentration of 1 µM at 5 hrs after moving cells to sporulation media (two replicates, each with 71 ≤ n ≤ 148 cells). *ndt80*Δ is strain ÜB27968, WT is strain ÜB21380, and *cdc28-as* is strain ÜB36314. **D.** Time-lapse microscopy of *ime2-as* cells treated with DMSO (left) or 1-NA-PP1 (right) to a final concentration of 20 µM at 5 hrs after moving cells to sporulation media. Cells contain reporter 2XmCherry-SV40NLS (pink) and fluorescently tagged histone Htb1-eGFP (green) in a mutant *ime2-as* background (strain ÜB25646). **E.** Percentage of cells that went through meiosis II (MII) with vNEBD initiation and barrier return (black), vNEBD initiation only (gray), or vNEBD initiation (white). Cells treated as in part D (45 ≤ n ≤ 118 cells). *IME2* is strain ÜB21380 and *ime2-as* is strain ÜB25646. **F.** Time-lapse microscopy of cells with reporter 2XmCherry- SV40NLS (pink) and fluorescently tagged histone Htb1-eGFP (green) in a mutant *ime2- st* background (strain ÜB27197). Cells with a constitutively active allele of Ime2 (*ime2- st*) under control of an inducible promoter (pGAL) were treated with either a vehicle (+EtOH) or the inducer (+ß-estradiol) in prophase arrest (*ndt80*Δ). For all images, the maximum-intensity z-projection is shown, and the scale bar is 2 µm. Full sample size information can be found in the Materials and Methods.

Besides regulating the canonical CDK, Ndt80 is also required for activating Ime2, a conserved, CDK-like kinase that is specifically expressed in meiotic cells. To block Ime2 activity during meiosis II, we utilized a conditional allele (*ime2-as*) that is sensitive to the chemical inhibitor 1-NM-PP1 (Benjamin et al., 2003), similar to our strategy for Cdc28. Excitingly, we found that vNEBD failed to occur when Ime2 activity was blocked (Fig. 5D-E). Since *ime2-as* mutants can undergo meiosis II chromosome segregation in the absence of vNEBD, our findings demonstrate that changes to nucleocytoplasmic transport can be uncoupled from nuclear divisions.

We next tested whether Ime2 was sufficient to initiate vNEBD. In cells that were arrested in meiotic prophase I (*ndt80*Δ), we induced expression of a constitutively active Ime2 (pGAL-*IME2st*; Sia and Mitchell 1995). vNEBD did not occur under these conditions (Fig. 5F), suggesting the involvement of additional factors besides Ime2 in regulating vNEBD. This finding is in contrast to other Ime2-dependent events in meiosis II, namely translational activation of a meiotic gene cluster and mitochondria detachment from the cell cortex, where Ime2 activity is both necessary and sufficient (Berchowitz et al. 2013; Sawyer et al. 2019). Altogether, our data conclusively link the initiation of vNEBD with regulation of the meiotic program via Ndt80 and Ime2, laying the groundwork for future identification of the drivers of nuclear permeability barrier disruption.

### vNEBD completion depends on the Glc7 phosphatase regulator Gip1

We next tried to gain mechanistic insight into the regulation of vNEBD completion. Our meiotic benchmarking experiments revealed that prospore membrane closure is tightly temporally correlated with the completion of vNEBD (Fig. 3E; on average 6.2 minutes before vNEBD ends). Thus, it is possible that gamete cytokinesis facilitates karyokinesis and, in turn, helps to efficiently end vNEBD. The prolonged vNEBD observed in *spo21*Δ mutants is consistent with this possibility (Fig. S4A). To more explicitly test a link between gamete cytokinesis and vNEBD completion, we utilized a mutant, *ama1*Δ, that disrupts prospore membrane closure. Ama1 is a meiosis-specific regulator of the anaphase promoting complex (APC) that contributes to PSM closure by targeting the leading edge complex for degradation (Diamond et al. 2009; Cooper et al. 2000). We found that even though *ama1*Δ cells successfully initiated vNEBD with similar timing as wild type, a noticeably smaller fraction completed vNEBD (Fig. 6A, C, S6A), exhibiting a significantly prolonged vNEBD duration (Fig. 6D). Therefore, it is possible that prospore membrane closure mediated by Ama1 is required for the timely and efficient completion of vNEBD.

**Figure 6.**
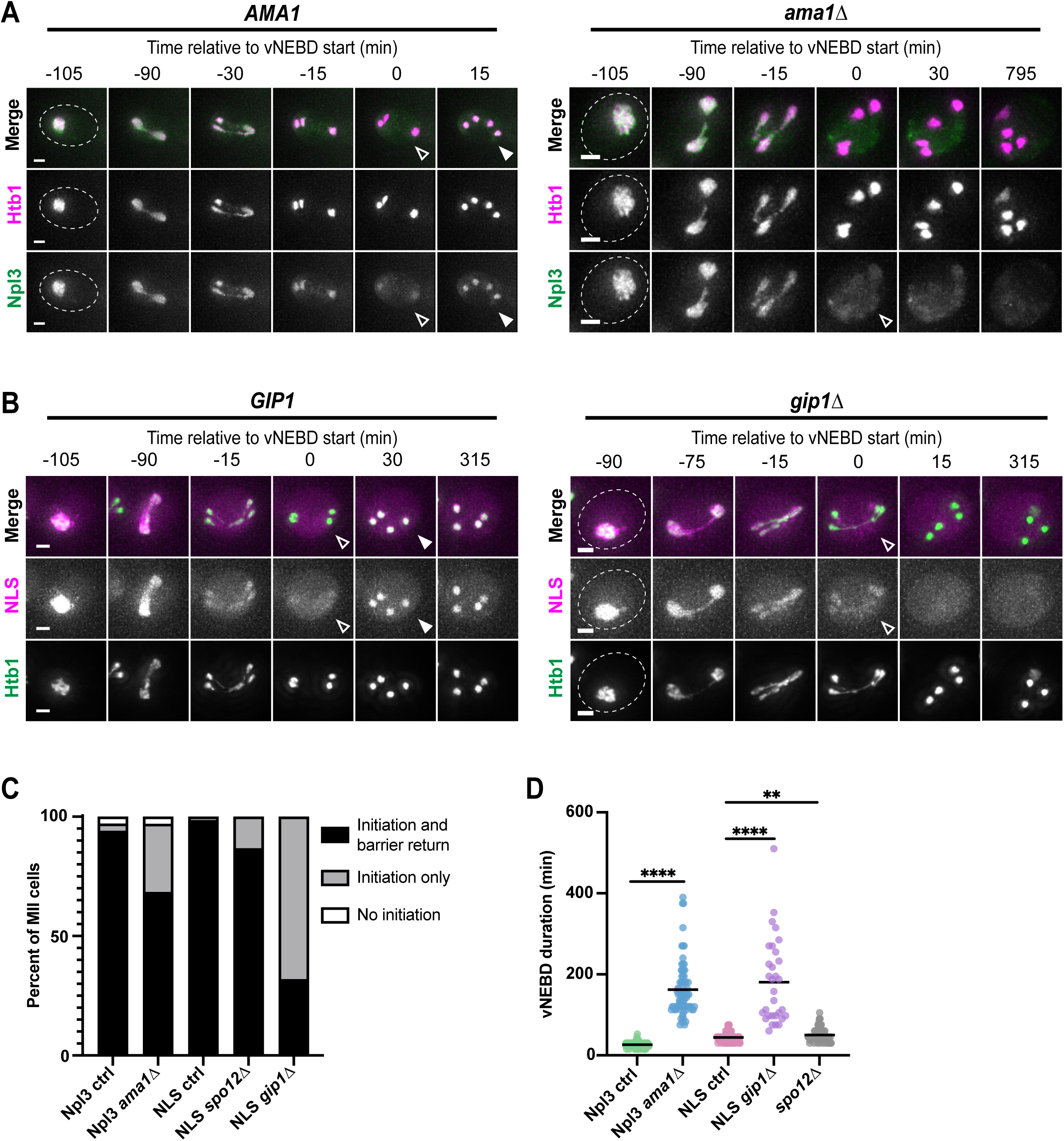
vNEBD completion depends on the Glc7 phosphatase regulator Gip1. A-B. Time-lapse microscopy of a cell progressing through meiosis. Open arrows indicate the start of vNEBD, and closed arrows represent the end of vNEBD. Maximum intensity projection shown and the scale bar is 2 µm. **A.** Cells contain fluorescently tagged histone Htb1-mCherry (pink) and nucleoplasmic protein GFP-Npl3 (green) in an *AMA1* background (left; strain ÜB18509) or a mutant *ama1*Δ background (right; strain ÜB20607). **B.** Cells contain reporter 2XmCherry-SV40NLS (pink) and fluorescently tagged histone Htb1-eGFP (green) in a *GIP1* background (left; strain ÜB21380) or a mutant *gip1*Δ background (right; strain ÜB21614). **C.** Percentage of cells that went through meiosis II (MII) with vNEBD initiation and barrier return (black), vNEBD initiation only (gray), or no vNEBD initiation (white) for various mutant backgrounds. Npl3 ctrl (strain ÜB18509) and Npl3 *ama1*Δ (strain ÜB20607) contain Htb1-mCherry and GFP- Npl3. NLS ctrl (strain ÜB21380), *spo12*Δ (strain ÜB36318), and *gip1*Δ (strain ÜB21614) all contain Htb1-eGFP and 2XmCherry-SV40NLS. Two replicates used for each strain (53 ≤ n ≤ 123 cells per replicate). **D.** Duration of vNEBD for various mutant backgrounds. Two replicates per strain with 53 ≤ n ≤ 123 cells per replicate. The strains are the same as in A. Lines represent the mean duration. Full sample size information can be found in the Materials and Methods.

Since the meiotic kinase Ime2 is required for vNEBD initiation, we next assessed whether phosphatases might play a role in vNEBD completion. The phosphatase Cdc14 resets the phosphorylation state of proteins targeted by Cyclin-CDK phosphorylation (Visintin, Hwang, and Amon 1999). Its activation by the Cdc14-early Release Factor (FEAR) response pathway is required for two consecutive chromosome divisions. In cells lacking *SPO12*, FEAR signaling is disrupted and, consequently, these cells undergo a single meiotic division akin to meiosis II, resulting in two spores (Marston, Lee, and Amon 2003). We monitored *spo12*Δ mutants to test whether the FEAR network, and thus Cdc14, is involved in vNEBD regulation. The vast majority (∼85%) of *spo12*Δ cells successfully initiated and completed vNEBD (Fig. 6C, S6B), with only a minor increase in vNEBD duration relative to wild type (Fig. 6D). FEAR signaling and Cdc14 are therefore dispensable for vNEBD. These findings further demonstrate that the complex nuclear morphology of the five-way division seen in wild-type cells can be uncoupled from vNEBD.

We finally interrogated whether vNEBD completion is regulated by Gip1, a meiosis-specific regulator of the serine-threonine PP1 phosphatase Glc7 that is upregulated during meiosis II in an Ime2-dependent manner (Tachikawa et al. 2001; Brar et al. 2012). Gip1 localization is dynamic throughout meiosis (Nakamura et al. 2017). In particular, Gip1 switches its localization from growing prospore membranes to gamete nuclei upon prospore membrane closure (Nakamura et al. 2017), coincident with vNEBD completion (Fig. 3E). Excitingly, we found that while *GIP1* was dispensable for vNEBD onset, *gip1*Δ mutants exhibited a prominent defect in vNEBD completion (Fig. 6B). Most *gip1*Δ cells never re-establish their nucleocytoplasmic permeability barrier (Fig. 6C), and those that do exhibit only weak re-accumulation of NLS-containing proteins in the nucleus with significant delays compared to wild-type cells (Fig. 6D). These findings indicate that PP1/Glc7 activity, controlled by Gip1, is required for vNEBD completion, leading to the restoration of nuclear and cytoplasmic compartments in gametes.

## Discussion

We have shown that meiosis II in the budding yeast *S. cerevisiae* features tightly regulated vNEBD, a process also observed in the distantly related *S. pombe* (Asakawa et al. 2010; Arai et al. 2010). vNEBD, previously thought to be unique to *S. pombe*, is characterized by semi-closed nuclear division, with bidirectional mixing between the cytoplasm and nucleoplasm (Fig. 1). This process occurs in the presence of an intact NE (King et al. 2019, Fig. S1F) and without the dispersal of channel nucleoporins (King et al. 2019, Fig. 3B). Remarkably, vNEBD diverges from all other known examples of semi-closed and semi-open divisions, which rely on either the dispersal of channel nucleoporins or the formation of holes in the NE via local NEBD (reviewed in Walsh, King and Ünal 2024). Our data show that vNEBD is not driven by the relocalization of RanGAP (Fig. 2) nor by any of the known NPC remodeling events during budding yeast meiosis (Fig. 4), leaving the mechanism of nuclear permeabilization an exciting mystery. This process is intricately coordinated with the meiotic program (Fig. 3), with vNEBD onset regulated by the transcription factor Ndt80 and the CDK-like kinase Ime2 (Fig. 5), while its resolution is controlled by the PP1 phosphatase regulatory subunit Gip1 (Fig. 6). Importantly, vNEBD highlights a significant distinction between nuclear divisions in mitosis and meiosis in budding yeast, alongside other differences such as NPC remodeling and sequestration. This finding contributes to a growing body of work demonstrating that nuclear remodeling is dynamically tailored to meet the distinct demands of different developmental stages within the same organism.

Interestingly, in contrast to other documented cases of semi-closed division, vNEBD exhibits incomplete diffusion of certain proteins between the nuclear and cytoplasmic compartments. For instance, endogenous nuclear proteins such as Npl3 and Pus1 remain partially enriched in the nucleus (Fig. 1A, S1B), while our NES reporter shows partial exclusion from the nucleus during vNEBD (Fig. 1E). Furthermore, large structures like ribosomes are robustly excluded from the nucleus during yeast vNEBD (Fig. S1G), consistent with the observed integrity of the NE (P. B. Moens 1971; Peter B. Moens and Rapport 1971; King et al. 2019). Altogether, these findings suggest that macromolecule movement has limitations during vNEBD, as opposed to the complete dispersal of nucleoplasmic proteins observed in organisms with semi-open (NE fenestration) or canonical semi-closed (partial NPC disassembly) divisions. For example, in *S. japonicus*, partial NEBD leads to rapid and complete dispersal of an endogenous nucleoplasmic protein (Nph6) within 10 seconds (Yam et al. 2011). Similarly, partial NPC disassembly during mitosis results in the complete dispersal of a synthetic NLS reporter in both *Dictyostelium discoideum* (Mitic et al. 2023) and *Aspergillus nidulans* (De Souza et al. 2004; Suresh et al. 2017). Interestingly, the partial retention or exclusion of individual proteins from the nucleus does not appear to be strictly size-dependent (compare Fig. 1A-B, S1A-B to S1C). For example, the larger RanGAP Rna1 (∼130 kDa with the 3x GFP tag) is only weakly excluded from the nucleus, while the smaller NES reporter (∼84 kDa with the 3x GFP tag) is more strongly excluded (compare Fig. 1E to Fig. 2C). Similarly, the duration of vNEBD does not correlate with protein size (Fig. S1C). We hypothesize that protein-protein interactions, including those involving transport regulators, might influence the extent and duration of protein relocalization during vNEBD.

vNEBD is tightly integrated into the meiotic program, occurring with very reproducible timing relative to other meiotic nuclear remodeling events (Fig. 3). However, even though these events are temporally correlated, we found that they do not drive vNEBD initiation or completion. Of particular note, NPC sequestration (*spo21*Δ) and nuclear basket return to nascent spore nuclei (*nup60*Δ and *nup60-*Δ*AH*) can be uncoupled from vNEBD (Fig. 4). Moreover, channel nucleoporins remain associated with the core NPC structure during meiosis (King et al. 2019), supporting the conclusion that vNEBD is not driven by known NPC remodeling mechanisms. The unique nuclear morphology of budding yeast anaphase II is also dispensable for vNEBD, as demonstrated by the fact that the *spo12*Δ mutant successfully executed vNEBD despite undergoing only a single meiotic division (Fig. 6A, S6B). Although GUNC formation still likely occurs during the single division in *spo12*Δ mutants (Fuchs and Loidl 2004; King et al. 2019), the apparent lack of GUNC formation in *S. pombe* suggests that it does not play a direct role in vNEBD (Asakawa et al. 2010; Arai et al. 2010). It remains possible, however, that novel and uncharacterized NPC remodeling events could be driving vNEBD. For example, although single NPC insertion events are not thought to disrupt nuclear integrity (Doucet and Hetzer 2010; Rothballer and Kutay 2013), the large-scale turnover of NPCs during meiosis II may necessitate a mass NPC biogenesis event that could temporarily disrupt nuclear compartmentalization.

We instead favor the hypothesis that meiotic nuclear remodeling events are temporally coordinated because they may share regulatory drivers (e.g., the master transcription factor Ndt80). We speculate that vNEBD – like many other cell-cycle coupled events– is driven by phosphorylation and dephosphorylation. This is supported by the requirement of meiosis-specific kinase Ime2 in initiating vNEBD (Fig. 5) and the meiosis-specific phosphatase regulator Gip1 in vNEBD completion (Fig. 6). Intriguingly, Ime2 (tested by induction of a constitutively active Ime2 allele, *ime2st*) does not appear to be sufficient for driving vNEBD initiation. This suggests the involvement of additional, unidentified regulators that are yet to be discovered. Future efforts should be directed towards identifying whether nuclear transport proteins are modified during meiosis II, either at the protein expression level or post-translationally by phosphorylation. Although localization of Rna1 (RanGAP) to the nucleus does not drive vNEBD (Fig. 2), modification of core directional transport machinery could explain a mass mixing event like vNEBD.

The remarkable conservation of vNEBD between *S. pombe* and *S. cerevisiae* – organisms that are diverged by over 400 million years of evolution (Sipiczki 2000) – suggests a conserved functional significance. Notably, no known mutants that prevent vNEBD in either *S. cerevisiae* or *S. pombe* result in viable spores (Fig. 5; Asakawa et al. 2010). In *S. cerevisiae*, perturbation of the meiotic CDK-like kinase Ime2 allows for a meiosis II division without vNEBD (Fig. 5E). In *S. pombe*, loss of Spo5, a meiosis-specific RNA-binding phosphoprotein that regulates CDK activity (Arata et al. 2014), results in a similar phenotype (Flor-Parra et al. 2018). In both systems, the pleiotropic roles of proteins identified as regulating vNEBD have precluded direct assessment of the contribution of vNEBD to gamete health, as neither mutant is capable of producing viable spores. More thorough characterization of vNEBD mechanisms will allow for its separation from other essential meiotic functions and assessment of its function.

vNEBD provides a unique mechanism by which cells can achieve the rapid and extensive mixing of proteins between cytoplasm and nucleoplasm. One potential role of vNEBD is timely breakdown of the meiosis II spindle, as has been demonstrated in *S. pombe* (Flor-Parra et al. 2018). Consistent with this notion, meiosis II spindle breakdown occurs directly following vNEBD in *S. cerevisiae* (Fig. 3A, D). We speculate, however, that additional meiosis-specific constraints might underlie the conserved disruption of nucleocytoplasmic compartmentalization during meiosis from yeast to humans. Mass exchange of proteins across compartments might facilitate degradation of harmful proteins and, in doing so, contribute to the rejuvenation of gametes. Further investigation of the mechanism and functional significance of vNEBD has the potential to reveal novel nuclear biology and quality control in meiosis.

## Materials and Methods

### Yeast strains, plasmids, and primers

All strains in this study are derived from SK1 and strain details can be found in Table S1. The following have been previously described: *HTB1-mCherry* (Matos et al. 2008); *ndt80*Δ (Xu et al. 1995); *GFP-NPL3* (Gilbert, Siebel, and Guthrie 2001); *2XmCherry- SV40NLS* (Monje-Casas and Amon 2009); *RPL26b-HTA-GFP* (Eisenberg et al. 2018); *PIL1-VH16* (Otto et al. 2021); *TUB1-GFP* (Straight et al. 1997); *CDC14-GFP* (Tomson et al. 2009); *NUP49-GFP*, *NUP60-GFP* (King et al. 2019); *nup60*Δ, *nup60-*Δ*AH*, (King et al. 2022); *CIT1-GFP, spo20(51-91)-yeGFP, spo21*Δ*, spo12*Δ*, ama1*Δ (Sawyer et al. 2019); *cdc28-as* (F88G; Bishop et al. 2000); *ime2-as* (M146G; Benjamin et al. 2003); *IME2st* (lacks the C-terminal 241 amino acids; Sia and Mitchell 1995).

Deletions and endogenous tags were made via established PCR methods (Janke et al. 2004; Longtine et al. 1998; Sheff and Thorn 2004). All primers used for strain construction are listed in Table S2, and all plasmids used are listed in Table S3. All single integration vectors were linearized with PmeI before integration into the genome.

The NES reporter was constructed via restriction cloning using the PKI NES from Shaikhqasem et al. 2020 fused to 3XeGFP and placed under the ARO10 promoter for strong meiotic expression in a TRP1 single integration vector. Following linearization with PmeI, the construct was integrated at the TRP1 locus. The nuclear envelope marker was constructed by fusing 2XeGFP to the NLS, linker, and transmembrane domain of Heh2 (293-378). The 2XmCherry-SV40NLS was a gift from the Sorger lab.

### Sporulation conditions

Sporulation was induced by starvation. Cells were first grown in YPD (1% yeast extract, 2% peptone, 2% glucose, 22.4 mg/L uracil, and 80 mg/L tryptophan) to saturation (OD600 ≥ 10) by incubating cells on room temperature shaker for approximately 24 hours. The culture was diluted to a concentration of OD600 = 0.2 in BYTA (1% yeast extract, 2% bacto tryptone, 1% potassium acetate, and 50 mM potassium phthalate) and grown on a shaker at 30°C overnight. When cultures reached OD600 ≥ 5, cells were pelleted and washed in sterile MilliQ water before resuspending in sporulation media (0.5% potassium acetate alone, 2% potassium acetate supplemented with amino acids (40 mg/L adenine, 40 mg/L uracil, 10 mg/L histidine, 10 mg/L leucine and 10 mg/L tryptophan) pH adjusted to 7 with acetic acid). Sporulation cultures were always set up at OD600 = 1.85 and incubated on a shaker at 30°C. For all culturing conditions, flasks were 10X the volume of media used to allow for proper aeration.

After 5 hours in sporulation media, *cdc28-as* cells were treated with 1-NM-PP1 to a final concentration of 1 µM, or the equivalent volume of DMSO, and *ime2-as* cells were treated with 1-NA-PP1 to a final concentration of 20 µM or the equivalent volume of DMSO. To induce *pGAL10-IME2st* expression, 1 µM of beta-estradiol or the equivalent volume of DMSO was added after 6 hours in sporulation media.

### Time lapse microscopy

All time lapse images were acquired on the DeltaVision Elite wide-field fluorescence microscope (GE Healthcare) using a 60x/1.42 NA oil-immersion objective. 8 z-slices were taken 1 µm apart. Images were deconvolved using the SoftWorx software (GE Healthcare). Images were either maximum-intensity z-projected or sum-projected using Fiji software. Imaging channels, laser intensity, plate type, and frequency of image acquisition are specified in Table S4.

Cells were imaged in either glass-bottom 96-well plates (Corning) or on a CellASIC ONIX Microfluidic Platform (EMD Millipore) using Y04E plates. During imaging, cells were kept at 30°C using an environmental chamber. We used conditioned sporulation media (filter sterilized after 5 hours of sporulation) to improve sporulation efficiency. For imaging using the 96-well plate, cells were harvested from shaking cultures and adhered to the bottom of the wells using concanavalin A, and 100 µl of conditioned sporulation meida was added. For the microfluidic CellASIC plates, cells were harvested from shaking cultures and loaded onto the imaging stage using 8 psi pressure for 5 s. Conditioned sporulation media flowed over cells at a rate of 2 psi for the rest of the image acquisition time.

To assess spore germination following asynchronous barrier return (differences in vNEBD duration), cells were imaged using the CellASIC system throughout meiosis (Table S4, Figure S2C) to track vNEBD dynamics. At approximately 24 hours post inoculation in sporulation media, the media flow was switched to YPD. Polarized light (32%T POL, 0.1s) images were acquired every 15 minutes for 8 hours with 8 z-slices, 1 µm apart.

### High resolution imaging

High resolution images were acquired using live cells on the Airyscan LSM 900 (Zeiss) using the Super Resolution (SR) Airyscan acquisition settings. Z-slices were taken 0.15 µm apart over the range set per cell. Images were acquired without averaging and at the maximum scan speed, switching channels every slice. Cells were harvested from shaking cultures and adhered to an 8-well dish using concanavalin A, and 150 µl of conditioned sporulation media was added. Cells were maintained at 30°C during imaging using an environmental chamber and heated stage. Specifics on imaging conditions per experiment can be found in Table S4.

### Image quantification

All images were processed in FIJI. To measure nuclear exclusion, line scans were drawn across each nucleus/nuclear lobe and the raw integrated density was measured across the line for a single z-slice in the approximate center of that nucleus, as assessed by histone Htb1 signal. Raw fluorescence values were then either plotted or normalized into a single metric as the lowest intensity measured across the line scan divided by the highest value. The average of this ratio was calculated per cell (each progenitor cell contains four gamete nuclei). We observe that nuclei that fail to successfully package (ex: fail to initiate PSM formation) often lose all nuclear integrity before dying. Because we only wanted to represent changes to nuclear integrity due to vNEBD, we omitted nuclei from our analysis if they had robust enrichment of cytoplasmic protein signal within the nucleus that differed from the other three nuclei in the cell. Cells were identified as being in late MII (“MII” in figure annotations) by the separation of chromatin into four lobes that are extended at opposite sides of the cell, often slightly oblong in shape, with no rDNA visible at the center of the division. Cells were identified as immature spores (“Spores” in figure annotations) by the distinct dark ring that separates the cytoplasmic signal in the spore from the rest of the progenitor cell (which we speculate to be the spore wall) but had not yet undergone ascal collapse. To measure whole-cell fluorescence, sum z-projections were generated in Fiji, and cell boundaries were gated using polarized light images as a reference. The raw integrated density was measured in both channels within the cell boundary for all time points, and data was normalized to the first time point of signal dispersal.

Nuclear barrier loss and return was only scored in cells that underwent MII, with the exception of *spo12*Δ cells, for which cells that underwent any genetic division were scored, and division incompetent mutants, for which all cells were scored including control cells that did not enter divisions.

For both timing experiments and to measure vNEBD duration, the timing of vNEBD initiation was assessed as the first time point that nuclear signal of a nucleoplasmic reporter decreased, and vNEBD end (barrier return) was scored as the first time point that signal of a nucleoplasmic reporter increased in the nucleus. End of anaphase II was scored as the point of maximum chromatin extension following rDNA segregation. Meiosis II spindle breakdown was assessed using exogenous Tub1-GFP to note the first time point that the spindle became fragmented, instead of a set of continuous lines. Mitochondrial collapse onto the nucleus was marked as the first time point that the mitochondria (marked with Cit1-GFP) localized away from the nuclear periphery (observed using polarized light images) and surrounded the nucleus (marked with histone Htb1). Nuclear basket return (marked with Nup60-GFP) is the first time point that the nuclear basket signal is enriched in gamete nuclei compared to the GUNC. Nuclear pore complex sequestration start (marked with Nup49-GFP) was scored as the first time point of asymmetric distribution of the NPC signal between the GUNC (enriched) and gamete nuclei. Cdc14 release from and re-accumulation to the nucleolus was assessed as the first time point that punctate Cdc14-GFP signal disappeared (release) and the first time point nuclear puncta reappeared (accumulation). PSM closure was assessed using reporter spo20(51-91)-GFP, which contains a PSM-specific transmembrane domain. PSM closure is coincident with the PSMs becoming circular (“rounding up”) rather than elongated (Diamond et al. 2009).

Detailed information about the sample size for each experiment can be found in Table S5. Statistical analysis was performed using Prism (GraphPad Software) using nonparametric, two-tailed Mann-Whitney tests.

## Acknowledgements

We thank Tianyao Xiao, Benjamin Styler, Naohiro Kuwayama, Maia Reyes, Silvan Spiri, Claudia Medrano, Tina Sing, and Cyrus Ruediger for their comments on this manuscript. We thank Gloria Brar, Doug Koshland, Rebecca Heald, Arash Komeili, and the entire Brar-Ünal lab for their thoughtful feedback and discussions. We thank David Drubin for the use of his Airyscan microscope and Leanna Owen for technical support with Airyscan imaging.

This work was supported by funds from the National Institutes of Health (R01AG071801) and Astera Institute to E. Ünal; National Science Foundation Graduate Research Fellowships (DGE1752814, DGE2146752) and a National Institutes of Health Traineeship (T32 GM007232) to G.A. King and M.E. Walsh. G.A. King is a Howard Hughes Medical Institute Fellow of the Damon Runyon Cancer Research Foundation (DRG-2500-2).

## Supplemental Figures

**Figure S1 A-B, D, F.** Time-lapse microscopy of cells progressing through meiosis. Open arrows indicate the start of vNEBD, and closed arrows represent the end of vNEBD. Maximum intensity projection shown. **A.** Cells contain fluorescently tagged histone Htb1-mCherry (pink) and nucleoplasmic protein Trz1-GFP (green) (strain ÜB20609). **B.** Cells contain fluorescently tagged histone Htb1-mCherry (pink) and nucleoplasmic protein Pus1-GFP (green) (strain ÜB24450). **C.** The molecular weight of fluorescently tagged nucleoplasmic proteins compared to the average duration of vNEBD per reporter. 2XmCherry-SV40NLS (pink), GFP-Npl3 (green), Trz1-GFP (blue), and Pus1-GFP (black). **D.** (Left) Cells contain nucleoplasmic reporter 2XmCherry-SV40NLS (pink) and fluorescently tagged GFP-Npl3 (green) (strain ÜB20617). (Right) Whole-cell fluorescence was normalized to the first time point of vNEBD. Points represent the mean for each timepoint and error bars represent standard deviation. n = 30 cells. **E.** Live Airyscan images of cells expressing both NES-3XeGFP (green) and 2XmCherry- SV40NLS (pink) (strain ÜB38941). Closed arrowheads indicate nuclei that are in-focus and have robust exclusion of the NES reporter in immature spores. Open arrowheads point to nuclei in focus in MII cells with dispersed NLS signal, which have higher nuclear NES signal. Single z-slice shown. **F.** Cells contain nucleoplasmic reporter 2XmCherry- SV40 NLS (pink) and a nuclear envelope (NE) marker (green; NLS, linker, and transmembrane domain from Heh2 fused to GFP) (strain ÜB38993). **G.** (Left) Representative live Airyscan images of sporulating cells expressing fluorescently-tagged histone Htb1-mCherry (pink) and ribosomal protein Rpl26b-GFP (green) (strain ÜB40128). The cell on the left contains immature spores (post MII but pre-ascal collapse) and the cell on the right is in late MII. Arrowheads represent nuclei in focus with robust exclusion of Rpl26b. Single z-slice shown. (Right) Quantification of nuclear Rpl26b-GFP signal in cells staged at late MII (n = 21 cells) or immature (pre-ascal collapse) spores (n = 20 cells). Line scans measured signal intensity across each nucleus, and an average of the ratio of minimum to maximum intensity was taken for each cell, such that each point represents one cell. Mann-Whitney test, not significant. Scale bar is 2 µm for all images shown. Full sample size information can be found in the Materials and Methods.

**Figure S2 A.** Line scans from the cells depicted in Figure 2C of raw fluorescence intensities (arbitrary units). Purple lines represent scans from each nucleus in the MII cell and teal lines represent scans from each nucleus in the cell with spores. Distance across the nucleus is measured in µm with the gray box representing the approximate nuclear boundary. **B.** The percentage of cells that underwent an MII division with synchronous or asynchronous barrier return for nuclei. Pil1+Nup159 tether refers to cells containing both the Pil1-nanobody and the Nup159-nanobody. **C.** Time-lapse microscopy of a cell progressing through meiosis. Open arrows indicate the start of vNEBD, and closed arrows represent the end of vNEBD (barrier return) for each nucleus. Cells contain nucleoplasmic reporter 2XmCherry-SV40NLS (pink), fluorescently tagged RanGAP Rna1-3XeGFP (green), and Pil1 and Nup159 both fused to an anti-GFP nanobody (strain ÜB20609). Maximum intensity projection shown. **D.** Quantification of the difference in barrier return timing for different nuclei from the same progenitor cell (two replicates each strain, 80 ≤ n ≤ 120 cells per replicate). The difference from the first nucleus with barrier return and the last nucleus with barrier return is shown. Lines represent the means (no tether = 0.2 min, Pil1-tether = 20.4 min, Pil1- and Nup159- tether = 34.5 min). **E.** Quantification of spore germination for cells with both the Pil1- nanobody and Nup159-nanobody (strain ÜB36287). Cells with synchronous vs asynchronous barrier return were compared (n = 66 cells). Full sample size information can be found in the Materials and Methods.

**Figure S3 A-B.** Time-lapse microscopy of cells progressing through meiosis. Open arrows indicate the start of vNEBD, and closed arrows represent the end of vNEBD. Maximum intensity projection shown. Scale bar is 2 µm. **A.** Cells contain reporter 2XmCherry- SV40NLS (pink) and fluorescently tagged mitochondrial marker Cit1-GFP (green) (strain ÜB44005). Asterisk marks the first time point of mitochondrial collapse onto the nucleus. **B.** Cells contain reporter 2XmCherry-SV40NLS (pink) and fluorescently tagged phosphatase Cdc14-GFP (green) (strain ÜB33512). One asterisk marks the post-meiosis I release of Cdc14, and two asterisks mark the re-accumulation of Cdc14 in the nucleolus.

**Figure S4 A.** Duration of vNEBD in various control and mutant backgrounds in minutes. Strains are the same as in Figure 4C. Two replicates used for each strain (62 ≤ n ≤ 118 cells per replicate). Bars represent the mean duration of vNEBD in minutes (NLS ctrl = 44.1 min, *nup60*Δ = 80.6 min, Npl3 ctrl = 30.6 min, *spo21*Δ = 44.6 min, *nup60-*Δ*AH* = 29.2 min, not significant). **B.** Timing of vNEBD in various mutant and WT backgrounds (same as in 4C) relative to anaphase II. Full sample size information can be found in the Materials and Methods.

**Figure S5 A.** Time-lapse microscopy of a *cdc28-as* cell treated with 1-NM-PP1 as in Figure 5B. Cell contains reporter 2XmCherry-SV40NLS (pink) and fluorescently tagged mitochondrial marker Cit1-eGFP (green) (strain ÜB44007). Open arrows indicate the start of vNEBD, and closed arrows represent the end of vNEBD. Asterisk marks the first time point of mitochondrial collapse onto the nucleus. **B**. The difference in time between mitochondrial collapse and barrier loss in *CDC28* cells (strain ÜB44005) or *cdc28-as* cells (strain ÜB44007) treated with either DMSO (vehicle) or 1-NM-PP1 (chemical inhibitor) (63 ≤ n ≤ 114 cells). **C.** (Left) Percentage of cells that went through meiosis II (MII) with vNEBD initiation and barrier return (black), vNEBD initiation only (gray), or no vNEBD initiation (white) for *CLB3* cells (strain ÜB21380) or *clb3*Δ cells (strain ÜB38349). (Right) Duration of vNEBD in minutes in *clb3*Δ cells compared to wild-type *CLB3* cells. Lines represent the mean duration (*CLB3* = 44.1 min, *clb3*Δ = 46.1 min). Mann-Whitney test, not significant. Full sample size information can be found in the Materials and Methods.

**Figure S6 A.** Time-lapse microscopy of a cell progressing through meiosis. Open arrows indicate the start of vNEBD and closed arrow represents the end of vNEBD. Maximum intensity projection shown and the scale bar is 2 µm. Cells contain reporter 2XmCherry- SV40NLS (pink) and fluorescently tagged histone Htb1-eGFP (green) in a mutant *spo12*Δ background (strain ÜB36318). **B.** vNEBD onset relative to the end of anaphase II. Npl3 ctrl (strain ÜB18509) and Npl3 *ama1*Δ (strain ÜB20607) contain Htb1-mCherry and GFP-Npl3. NLS ctrl (strain ÜB21380), *spo12*Δ (strain ÜB36318), and *gip1*Δ (strain ÜB21614) all contain Htb1-eGFP and 2XmCherry-SV40NLS. Full sample size information can be found in the Materials and Methods.

