## Supplementary figures and images for "A Conserved Disruption of the Nuclear Permeability Barrier in Meiosis is Controlled by a Kinase-Phosphatase Pair in *Saccharomyces cerevisiae*"

### Supplemental Figure 1

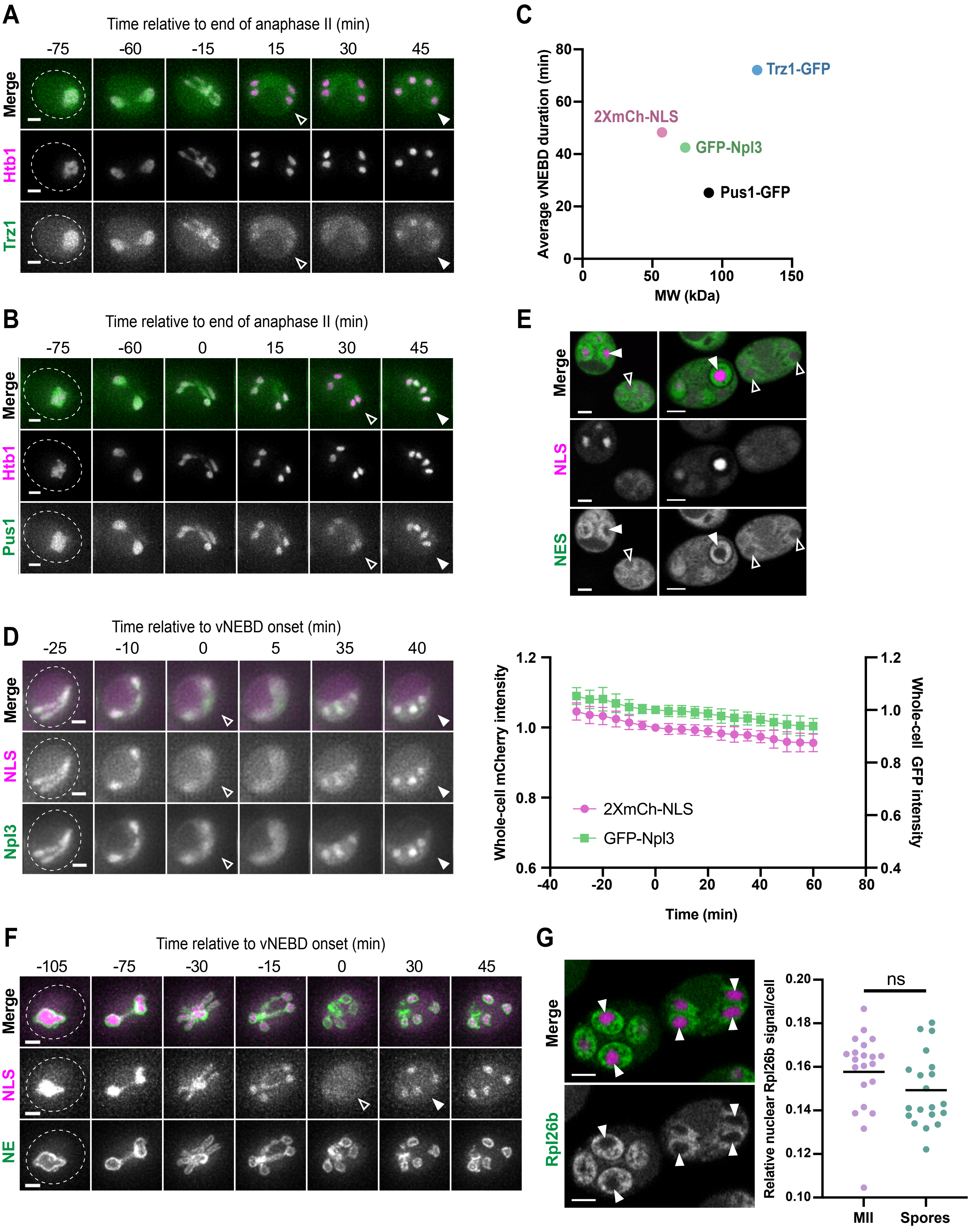

### Supplemental Figure 2

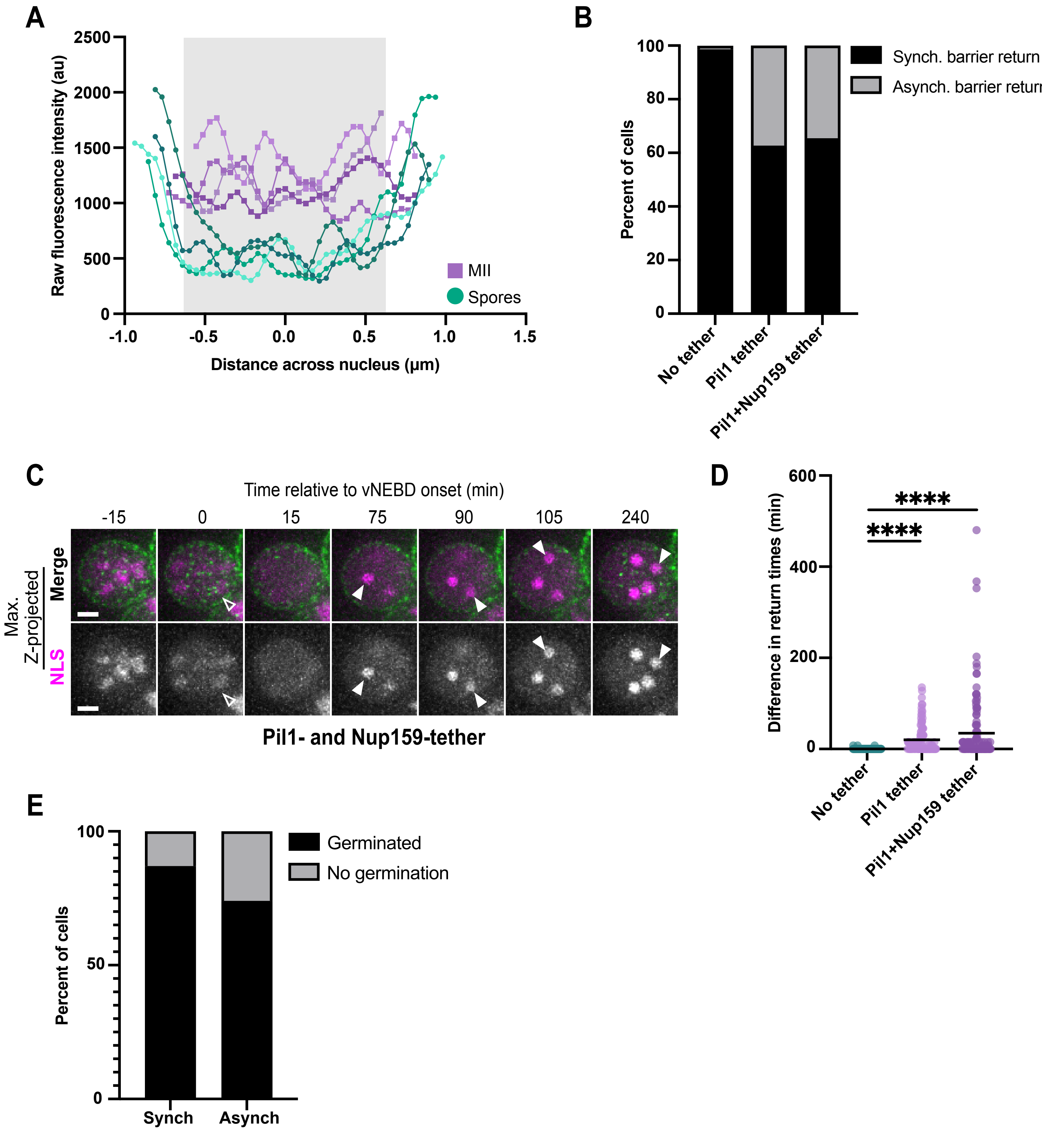

### Supplemental Figure 3

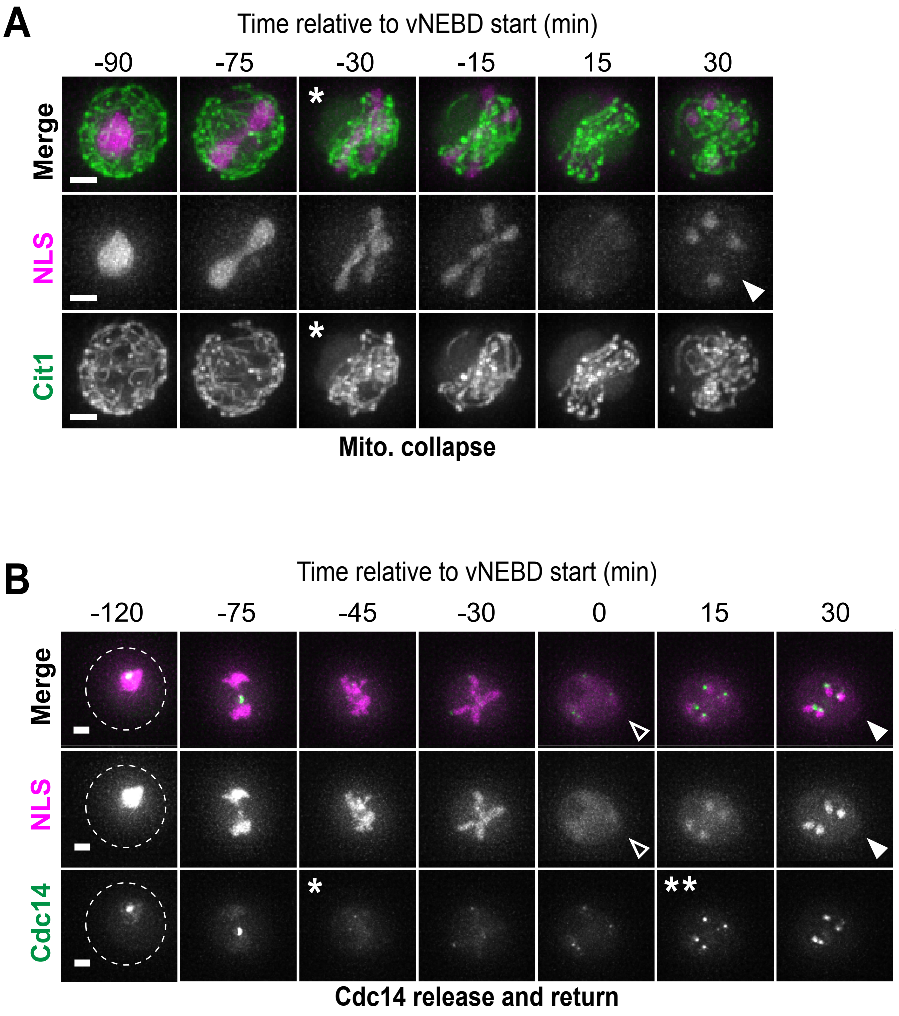

### Supplemental Figure 4

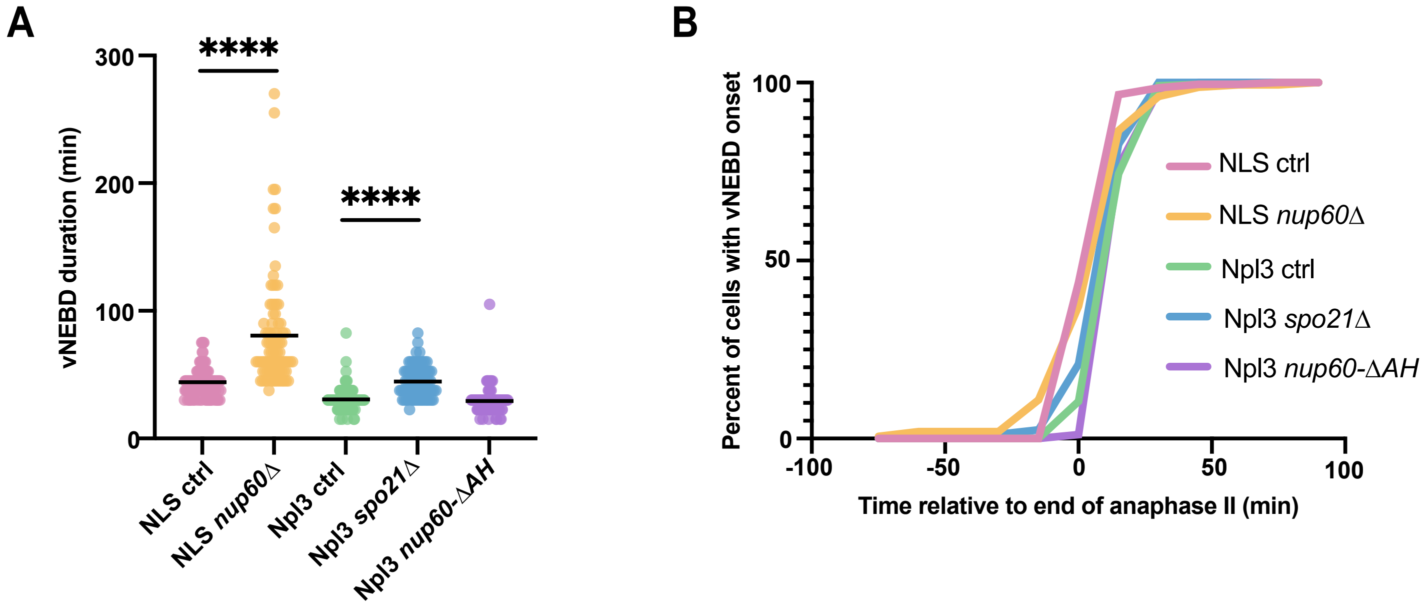

### Supplemental Figure 5

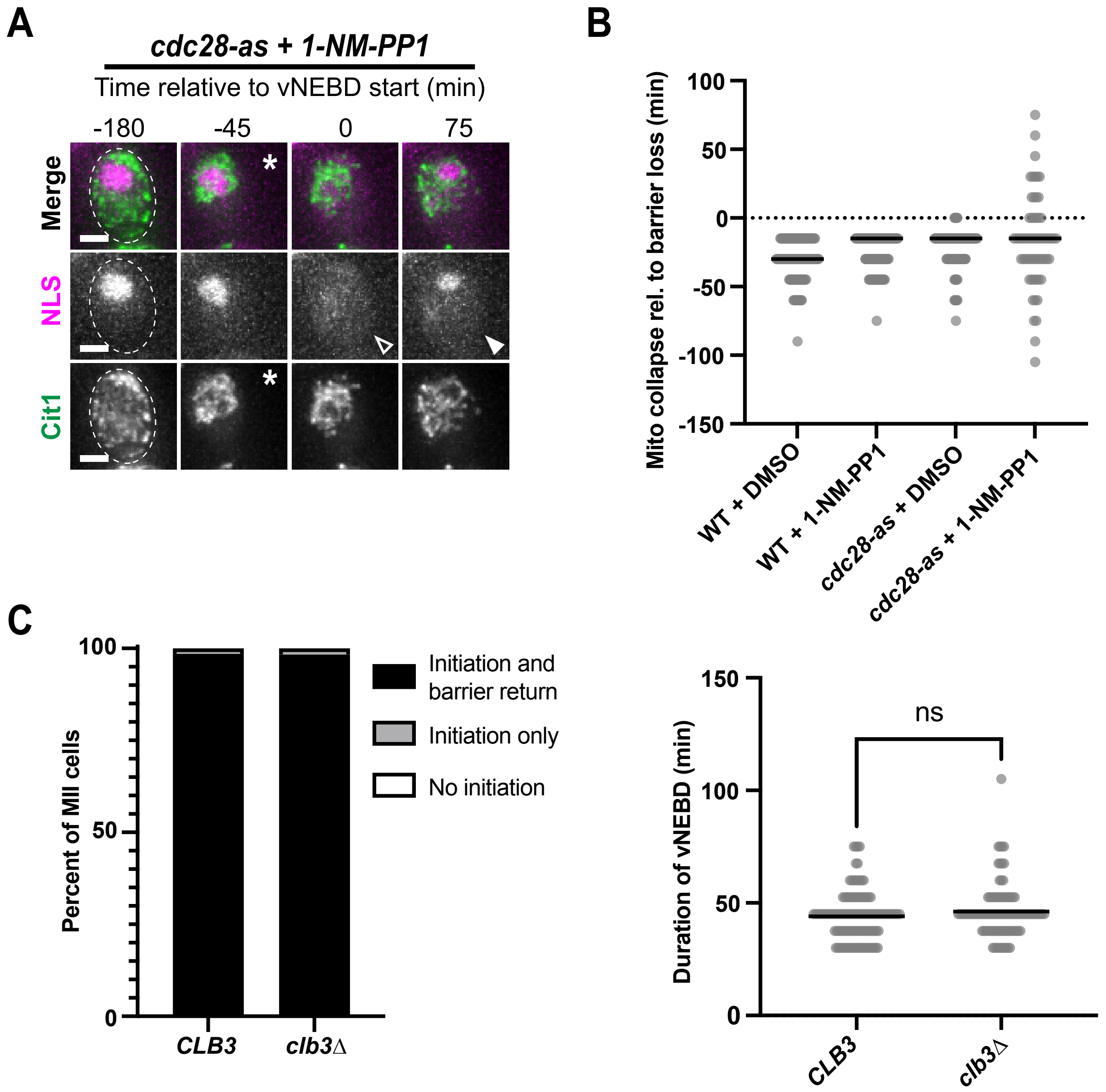

### Supplemental Figure 6

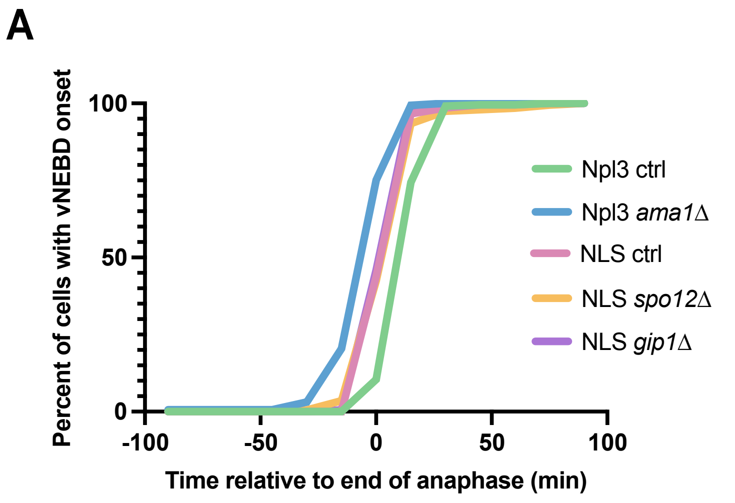

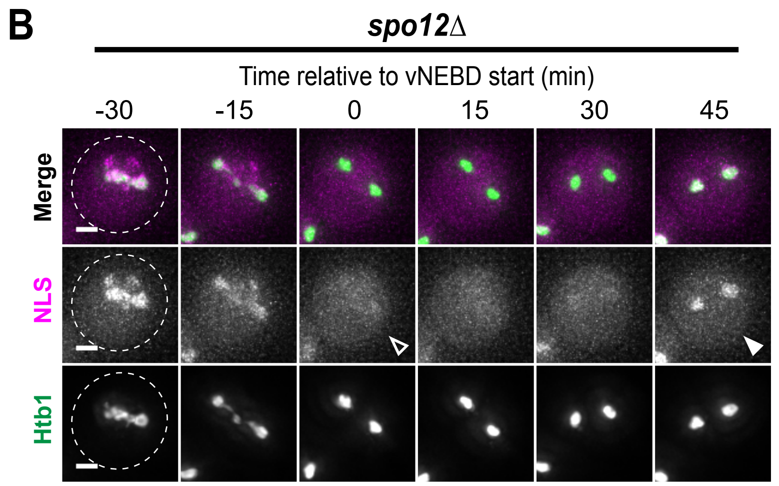
