## Supplemental Table 1 for "A Conserved Disruption of the Nuclear Permeability Barrier in Meiosis is Controlled by a Kinase-Phosphatase Pair in *Saccharomyces cerevisiae*"

**Table S1. Yeast strains used in this study.**

| **Strain** | **Genotype** |
| --- | --- |
| SK1  wild-type | *ho::LYS2 lys2 ura3 leu2::hisG his3::hisG trp1::hisG* |
| ÜB18509 | MATa/MATalpha his3::GFP-Npl3::HIS3/his3::GFP-Npl3::HIS3 HTB1-mCherry-HISMX6/HTB1-mCherry-HISMX6 |
| ÜB18513 | MATa/MATalpha Leu2::2xmCherry-SV40NLS::LEU2/Leu2::2xmCherry-SV40NLS::LEU2 Nup49-GFP::KanMX/Nup49-GFP::KanMX |
| ÜB20153 | MATa/MATalpha Rna1-3xEGFP::KanMX/Rna1-3xEGFP::KanMX HTB1-mCherry-HISMX6/HTB1-mCherry-HISMX6 |
| ÜB20155 | MATa/MATalpha Rna1-3xEGFP::KanMX/Rna1-3xEGFP::KanMX Leu2::2xmCherry-SV40NLS::LEU2/Leu2::2xmCherry-SV40NLS::LEU2 |
| ÜB20161 | MATa/MATalpha Prp20-GFP-HisMX/Prp20-GFP-HisMX HTB1-mCherry-HISMX6/HTB1-mCherry-HISMX6 |
| ÜB20607 | MATa/MATalpha ama1::KanMX6/ama1::KanMX6 his3::GFP-Npl3::HIS3/his3::GFP-Npl3::HIS3 HTB1-mCherry-HISMX6/HTB1-mCherry-HISMX6 |
| ÜB20609 | MATa/MATalpha trp1::TRZ1-WT-yEGFP::TRP1/trp1::TRZ1-WT-yEGFP::TRP1 HTB1-mCherry-HISMX6/HTB1-mCherry-HISMX6 |
| ÜB20617 | MATa/MATalpha his3::GFP-Npl3::HIS3/his3::GFP-Npl3::HIS3 Leu2::2xmCherry-SV40NLS::LEU2/Leu2::2xmCherry-SV40NLS::LEU2 |
| ÜB21380 | MATa/MATalpha Htb1-EGFP-KanMX/Htb1-EGFP-KanMX Leu2::2xmCherry-SV40NLS::LEU2/Leu2::2xmCherry-SV40NLS::LEU2 |
| ÜB21612 | MATa/MATalpha spo21::HygB/spo21::HygB Htb1-EGFP-KanMX/Htb1-EGFP-KanMX Leu2::2xmCherry-SV40NLS::LEU2/Leu2::2xmCherry-SV40NLS::LEU2 |
| ÜB21614 | MATa/MATalpha Gip1::NatMX/Gip1::NatMX Htb1-EGFP-KanMX/Htb1-EGFP-KanMX Leu2::2xmCherry-SV40NLS::LEU2/Leu2::2xmCherry-SV40NLS::LEU2 |
| ÜB24450 | MATa/MATalpha HTB1-mCherry-HISMX6/HTB1-mCherry-HISMX6 Pus1-GFP:KanMX/Pus1-GFP:KanMX |
| ÜB25072 | MATa/MATalpha Nup60::HygMX/Nup60::HygMX Htb1-EGFP-KanMX Leu2::2xmCherry-SV40NLS::LEU2/Leu2::2xmCherry-SV40NLS::LEU2 |
| ÜB25646 | MATa/MATalpha ime2-as1/ime2-as1 Htb1-EGFP-KanMX/Htb1-EGFP-KanMX Leu2::2xmCherry-SV40NLS::LEU2/Leu2::2xmCherry-SV40NLS::LEU2 |
| ÜB25839 | MATa/MATalpha Nup60-delta2-47-NatMX/Nup60-delta2-47-NatMX his3::GFP-Npl3::HIS3/his3::GFP-Npl3::HIS3 HTB1-mCherry-HISMX6/HTB1-mCherry-HISMX6 |
| ÜB27197 | MATa/MATalpha ndt80::LEU2/ndt80::LEU2 ura3::pGPD1-GAL4(848).ER::URA3/ura3::pGPD1-GAL4(848).ER::URA3 KanMX-pGAL-IME2st his3::GFP-Npl3::HIS3/his3::GFP-Npl3::HIS3 HTB1-mCherry-HISMX6/HTB1-mCherry-HISMX6 |
| ÜB27968 | MATa/MATalpha ndt80::LEU2/ndt80::LEU2 his3::GFP-Npl3::HIS3/his3::GFP-Npl3::HIS3 HTB1-mCherry-HISMX6/HTB1-mCherry-HISMX6 |
| ÜB33507 | MATa/MATalpha ura3::pTUB1-GFP-TUB1::URA3 Leu2::2xmCherry-SV40NLS::LEU2/Leu2::2xmCherry-SV40NLS::LEU2 |
| ÜB33512 | MATa/MATalpha CDC14-GFP-LEU2/CDC14-GFP-LEU2 Leu2::2xmCherry-SV40NLS::LEU2/Leu2::2xmCherry-SV40NLS::LEU2 |
| ÜB34583 | MATa/MATalpha his3::pATG8-link-yEGFP-SPO20(51-91)::HIS3/his3::pATG8-link-yEGFP-SPO20(51-91)::HIS3 Leu2::2xmCherry-SV40NLS::LEU2/Leu2::2xmCherry-SV40NLS::LEU2 |
| ÜB36206 | MATa/MATalpha Leu2::2xmCherry-SV40NLS::LEU2/Leu2::2xmCherry-SV40NLS::LEU2 Rna1-3xEGFP::KanMX/Rna1-3xEGFP::KanMX Pil1-antiGFP::URA3/Pil1-antiGFP::URA3 |
| ÜB36287 | MATa/MATalpha Leu2::2xmCherry-SV40NLS::LEU2/Leu2::2xmCherry-SV40NLS::LEU2 Rna1-3xEGFP::KanMX/Rna1-3xEGFP::KanMX Nup159-VH16::Ura3/Nup159-VH16::Ura3 Pil1-antiGFP::URA3/Pil1-antiGFP::URA3 |
| ÜB36314 | MATa/MATalpha Htb1-EGFP-KanMX/Htb1-EGFP-KanMX Leu2::2xmCherry-SV40NLS::LEU2/Leu2::2xmCherry-SV40NLS::LEU2 cdc28-as1 (F88G)/cdc28-as1 (F88G) |
| ÜB36318 | MATa/MATalpha Htb1-EGFP-KanMX/Htb1-EGFP-KanMX Leu2::2xmCherry-SV40NLS::LEU2/Leu2::2xmCherry-SV40NLS::LEU2 spo12::KanMX/spo12::KanMX |
| ÜB38349 | MATa/MATalpha clb3∆::HygMX/clb3∆::HygMX Htb1-EGFP-KanMX/Htb1-EGFP-KanMX Leu2::2xmCherry-SV40NLS::LEU2/Leu2::2xmCherry-SV40NLS::LEU2 |
| ÜB38411 | MATa/MATalpha trp1::pARO10-PKI-NES-3XGFP::TRP1/trp1::pARO10-PKI-NES-3XGFP::TRP1 HTB1-mCherry-HISMX6/HTB1-mCherry-HISMX6 |
| ÜB38941 | MATa/MATalpha Leu2::2xmCherry-SV40NLS::LEU2/Leu2::2xmCherry-SV40NLS::LEU2 trp1::pARO10-PKI-NES-3XGFP::TRP1/trp1::pARO10-PKI-NES-3XGFP::TRP1 |
| ÜB38993 | MATa/MATalpha leu2::pATG8-2xGFP-h2NLS-L-TM::LEU2/leu2::pATG8-2xGFP-h2NLS-L-TM::LEU2 trp1::pARO10-2XmCherry-SV40NLS::TRP1/trp1::pARO10-2XmCherry-SV40NLS::TRP1 |
| ÜB40128 | MATa/MATalpha rpl26b:Rpl26b-HTA-GFP-Kan/rpl26b:Rpl26b-HTA-GFP-Kan HTB1-mCherry-HISMX6/HTB1-mCherry-HISMX6 |
| ÜB44005 | MATa/MATalpha Leu2::2xmCherry-SV40NLS::LEU2/Leu2::2xmCherry-SV40NLS::LEU2 CIT1-GFP::His3MX6/CIT1-GFP::His3MX6 |
| ÜB44007 | MATa/MATalpha CIT1-GFP::His3MX6/CIT1-GFP::His3MX6 Leu2::2xmCherry-SV40NLS::LEU2/Leu2::2xmCherry-SV40NLS::LEU2 cdc28-as1 (F88G)/cdc28-as1 (F88G) |
| ÜB44905 | MATa/MATalpha Nup60-GFP-KanMX/Nup60-GFP-KanMX Leu2::2xmCherry-SV40NLS::LEU2/Leu2::2xmCherry-SV40NLS::LEU2 |
