## Supplemental Table 2 for "A Conserved Disruption of the Nuclear Permeability Barrier in Meiosis is Controlled by a Kinase-Phosphatase Pair in *Saccharomyces cerevisiae*"

Table S2. Primers used for strain construction.

| **Construct name** | **Forward Primer** | **Reverse Primer** |
| --- | --- | --- |
| Htb1-eGFP-KanMX | TACTAGAGCTGTTACCAAGTACTCTTCCTCTACTCAAGCACGTACGCTGCAGGTCGAC | TAAATAATAATATTAATTATAACCAAAGGAAGTGATTTCAATCGATGAATTCGAGCTCG |
| Pus1-GFP | gccttagaaattaaagttggtaagaaagaaggaaagggcaactcgatcaattcgagctcg | cagagcaacgaacctgaagtacaaccggaagcggcagctaatggtgacggtgctggttta |
| Rna1-3XeGFP | AGATGATCTTGCTGAACGTTTAGCTGAAACTGAAATCAAACGGATCCCCGGGTTAATTAA | GTCCACAGTTGATTGTGTTTATTTTTACTTTTATTCATAGGAATTCGAGCTCGTTTAAAC |
| Prp20-GFP | TGATGAGGACGCAGAAAAGAGAGCGGATGAAATGGATGATCGGATCCCCGGGTTAATTAA | ATTATGTCGATTTTCTTTTATTTATCTTTGTACTACTACCGAATTCGAGCTCGTTTAAAC |
| Nup159-VH16 | GCAAATTGGTGATTTCTTCAAAAATTTGAACATGGCAAAAggtgacggtgctggttta | TTATTAACGGCACTAACAACGTACATATAGCTAAATATCAtcgatgaattcgagctcg |
| clb3∆ | CATAGAGATATTTTGTCCTTTTTATCATTGCTTATTATAAcgtacgctgcaggtcgac | CTTTTTCCTTTGTTGATGCCATGTCTCGAGCTGAGGCTTTatcgatgaattcgagctcg |
| Gip1∆::NatMX | CAAATTTTTTGGTGAAGAGCTATTCAATTATTAGCTAATTCGGATCCCCGGGTTAATTAA | AATTTTACAGAGTATTGGCAGTTAAGTGTTGTTTTTTCGCGAATTCGAGCTCGTTTAAAC |
