## Supplemental Table 3 for "A Conserved Disruption of the Nuclear Permeability Barrier in Meiosis is Controlled by a Kinase-Phosphatase Pair in *Saccharomyces cerevisiae*"

Table S3. Plasmids used for strain construction.

| **Plasmid name** | **Description** | **Type** |
| --- | --- | --- |
| pÜB1 | pFA6a-kanMX6 | For endogenous tagging |
| ÜB6 | pFA6a-GFP(S65T)-His3MX6 | For endogenous tagging |
| pÜB76 | pKT127 (pFA6a-link-yEGFP-Kan) | For endogenous tagging |
| pÜB153 | pfa6a-NatMX4 | For endogenous tagging |
| pÜB182 | pfa6a-link-yoEGFP-Kan | For endogenous tagging |
| pÜB217 | pFA6a-hphNT1 | For endogenous tagging |
| pÜB233 | pYM27-eGFP-KanMX | For endogenous tagging |
| pÜB984 | pARO10-2xmCherry-SV40NLS | Single integration at the LEU2 locus |
| ÜB985 | pFA6-3xeGFP-KanMX6 | For endogenous tagging |
| pÜB1194 | pAE47-TRZ1-WT-yEGFP | Single integration at the TRP1 locus |
| ÜB1707 | pFA6a–link–VH16–CaURA3 | For endogenous tagging |
| pÜB2546 | pCR44_pLC605-pATG8-(2x)yeGFP-h2NLS-L-TM | Single integration at the LEU2 locus |
| pÜB2600 | pARO10-PKI-NES-3XGFP | Single integration at the TRP1 locus |
| pÜB2660 | pARO10-2XmCh-NLS | Single integration at the TRP1 locus |
