## Supplemental Table 4 for "A Conserved Disruption of the Nuclear Permeability Barrier in Meiosis is Controlled by a Kinase-Phosphatase Pair in *Saccharomyces cerevisiae*"

**Table S4. Imaging conditions.**

| **Live time-lapse widefield imaging** | | | | | | |
| --- | --- | --- | --- | --- | --- | --- |
| **Figure** | **Plate** | **RFP** | **GFP** | **POL** | **Z-sectioning** | **Time resolution** |
| 1A, S1A, S1B, 2B, 4B, 4E, 5A, 5F, 6A | Cell asics YO4E | 10%T mCherry 0.025s  EX: 575/25  EM: 632/60 | 10%T FITC  0.025s  EX: 475/28  EM: 523/36 | 32%T  0.1s | 8 slices, 1 µm apart | Image every 15 min. for 18 hrs |
| 1B, S1F, 2E, 2F, S2C, 3B, 3C, 3D, 3E, S3A, S3B, 4D, 6B, S6A | Cell asics YO4E | 32%T mCherry 0.025s  EX: 575/25  EM: 632/60 | 10%T FITC  0.025s  EX: 475/28  EM: 523/36 | 32%T  0.1s | 8 slices, 1 µm apart | Image every 15 min. for 18 hrs |
| 5B, 5D, S5A | 96-well plate | 32%T mCherry 0.025s  EX: 575/25  EM: 632/60 | 10%T FITC  0.025s  EX: 475/28  EM: 523/36 | 32%T  0.1s | 8 slices, 1 µm apart | Image every 15 min. for 18 hrs |
| S1D | Cell asics YO4E | 32%T mCherry 0.025s  EX: 575/25  EM: 632/60 | 10%T FITC  0.025s  EX: 475/28  EM: 523/36 | 32%T  0.1s | 8 slices, 1 µm apart | Image every 5 min. for 18 hrs |
| **Live SR Airyscan imaging** | | | | | | |
| **Figure** | **Plate** | **RFP** | **GFP** | **POL** | **Z-sectioning** | **Time resolution** |
| 1E | 8-well Lab-Tek dish | 0.35%  mCherry | 0.6%  eGFP | -- | Z slices across range of the cell, 0.15 µm apart | Single image |
| 2C | 8-well Lab-Tek dish | 0.35%  mCherry | 3%  eGFP | -- | Z slices across range of the cell, 0.15 µm apart | Single image |
| S1G | 8-well Lab-Tek dish | 0.35%  mCherry | 1.5%  eGFP | -- | Z slices across range of the cell, 0.15 µm apart | Single image |
| S1E | 8-well Lab-Tek dish | 2%  mCherry | 0.6%  eGFP | -- | Z slices across range of the cell, 0.15 µm apart | Single image |
