## Supplemental Table 5 for "A Conserved Disruption of the Nuclear Permeability Barrier in Meiosis is Controlled by a Kinase-Phosphatase Pair in *Saccharomyces cerevisiae*"

Table S5. Sample sizes.

| **Experiment** | **Relevant Figure(s)** | **Sample** | **Strain number** | **Sample size (n)** |
| --- | --- | --- | --- | --- |
| Tracking protein dispersal from the nucleus relative to meiotic divisions | Figure 1C, 1D, 3A | GFP-Npl3 | ÜB18509 | *Rep 1*: 119 cells  *Rep 2*: 112 cells |
|  |  | Trz1-GFP | ÜB20609 | *Rep 1*: 97 cells  *Rep 2*: 103 cells |
|  |  | Pus1-GFP | ÜB24450 | *Rep 1*: 65 cells  *Rep 2*: 106 cells |
|  |  | 2XmCherry-SV40NLS | ÜB21380 | *Rep 1*: 140 cells  *Rep 2*: 118 cells |
| Quantification of NES-3XGFP signal in the nucleus in meiosis using high resolution imaging (Airyscan SR) | Figure 1E | Cells in meiosis II | ÜB38411 | *Rep 1*: 14 cells  *Rep 2*: 28 cells |
|  |  | Pre-mature spores (post-meiosis II) | ÜB38411 | *Rep 1*: 27 cells  *Rep 2*: 18 cells |
| Quantification of whole-cell fluorescence through meiosis | Figure S1D | GFP-Npl3 2XmCherry-SV40NLS | ÜB20617 | 30 cells |
| Quantification of Rpl26b-GFP signal in the nucleus in meiosis using high resolution imaging (Airyscan SR) | Figure S1G | Cells in meiosis II | ÜB40128 | 21 cells |
|  |  | Pre-mature spores (post-meiosis II) | ÜB40128 | 20 cells |
| Quantification of RanGAP signal in the nucleus in meiosis using high resolution imaging (Airyscan SR) | Figure 2C | Cells in meiosis II | ÜB20153 | *Rep 1*: 13 cells  *Rep 2*: 20 cells |
|  |  | Pre-mature spores (post-meiosis II) | ÜB20153 | *Rep 1*: 8 cells  *Rep 2*: 16 cells |
| Measuring vNEBD onset time and duration in RanGAP tethered and untethered cells | Figure 2G, 2H, S2B, S2D | No tether | ÜB20155 | *Rep 1*: 108 cells  *Rep 2*: 106 cells |
|  |  | Pil1-tether | ÜB36206 | *Rep 1*: 79 cells  *Rep 2*: 106 cells |
|  |  | Pil1- and Nup159-tether | ÜB36287 | *Rep 1*: 103 cells  *Rep 2*: 120 cells |
| Germination of cells with asynchronous barrier return | Figure S2E | Pil1- and Nup159-tether | ÜB36287 | 66 cells |
| vNEBD timing relative to NPC sequestration | Figure 3A, 3F | Nup49-GFP  2XmCherry-SV40NLS | ÜB18513 | *Rep 1*: 112 cells |
| vNEBD timing relative to mitochondrial collapse onto the nucleus | Figure 3A, 3F | Cit1-GFP  2XmCherry-SV40NLS | ÜB44005 | *Rep 1*: 107 cells  *Rep 2*: 116 cells |
| vNEBD timing relative to Cdc14 release from the nucleolus and re-accumulation | Figure 3A, 3F | Cdc14-GFP  2XmCherry-SV40NLS | ÜB33512 | *Rep 1*: 117 cells  *Rep 2*: 88 cells |
| vNEBD timing relative to nuclear basket return to gamete nuclei | Figure 3A, 3F | Nup60-GFP  2XmCherry-SV40NLS | ÜB44905 | *Rep 1*: 115 cells  *Rep 2*: 108 cells |
| vNEBD timing relative to MII spindle breakdown | Figure 3A, 3F | Tub1-GFP  2XmCherry-SV40NLS | ÜB33507 | *Rep 1*: 81 cells  *Rep 2*: 154 cells |
| vNEBD timing relative to PSM closure | Figure 3A, 3F | Spo20(51-91)-eGFP  2XmCherry-SV40NLS | ÜB34583 | *Rep 1*: 118 cells  *Rep 2*: 137 cells |
| Measuring vNEBD onset time and duration in NPC remodeling mutants | Figure 4C, S4A, S4B | GFP-Npl3 control | ÜB18509 | *Rep 1*: 112 cells  *Rep 2*: 117 cells |
|  |  | GFP-Npl3  *spo21∆* | ÜB21612 | *Rep 1*: 74 cells  *Rep 2*: 85 cells |
|  |  | GFP-Npl3  *nup60-∆AH* | ÜB25839 | *Rep 1*: 73 cells  *Rep 2*: 101 cells |
|  |  | 2XmCherry-SV40NLS control | ÜB21380 | *Rep 1*: 118 cells  *Rep 2*: 85 cells |
|  |  | 2XmCherry-SV40NLS  *nup60∆* | ÜB25072 | *Rep 1*: 85 cells  *Rep 2*: 62 cells |
| Measuring vNEBD penetrance and/or onset time various meiotic progression mutant backgrounds | Figure 5C | *ndt80∆* | ÜB27968 | *Rep 1*: 112 cells |
|  |  | *CDC28* +DMSO | ÜB21380 | *Rep 1*: 122 cells  *Rep 2*: 148 cells |
|  |  | *CDC28*  +1-NM-PP1 | ÜB21380 | *Rep 1*: 107 cells  *Rep 2*: 126 cells |
|  |  | *cdc28-as*  +DMSO | ÜB36314 | *Rep 1*: 106 cells  *Rep 2*: 145 cells |
|  |  | *cdc28-as*  +1-NM-PP1 | ÜB36314 | *Rep 1*: 124 cells  *Rep 2*: 147 cells |
|  | Figure 5E | *IME2*  +DMSO | ÜB21380 | 118 cells |
|  |  | *IME2*  +1-NA-PP1 | ÜB21380 | 115 cells |
|  |  | *ime2-as*  +DMSO | ÜB25646 | 110 cells |
|  |  | *ime2-as*  +1-NA-PP1 | ÜB25646 | 45 cells  (only cells that complete MII were included; treatment decreases MII completion) |
|  | Figure S5C | *clb3∆* | ÜB38349 | *Rep 1*: 82 cells  *Rep 2*: 104 cells |
|  | Figure 6C, S6A | *spo12∆* | ÜB36318 | *Rep 1*: 113 cells  *Rep 2*: 90 cells |
|  |  | *ama1∆* | ÜB20607 | *Rep 1*: 107 cells  *Rep 2*: 53 cells |
|  |  | *gip1∆* | ÜB21614 | *Rep 1*: 53 cells  *Rep 2*: 119 cells |
| Measuring vNEBD duration in various meiotic progression mutant backgrounds | Figure S5C | *clb3∆* | ÜB38349 | *Rep 1*: 104 cells  *Rep 2*: 79 cells |
|  | Figure 6D | *spo12∆* | ÜB36318 | *Rep 1*: 97 cells  *Rep 2*: 79 cells |
|  |  | *ama1∆* | ÜB20607 | *Rep 1*: 75 cells  *Rep 2*: 38 cells |
|  |  | *gip1∆* | ÜB21614 | *Rep 1*: 24 cells  *Rep 2*: 31 cells |
| Difference in mitochondrial collapse time and vNEBD onset | Figure S5B | *CDC28* +DMSO | ÜB44005 | 114 cells |
|  |  | *CDC28*  +1-NM-PP1 | ÜB44005 | 102 |
|  |  | *cdc28-as*  +DMSO | ÜB44007 | 104 |
|  |  | *cdc28-as*  +1-NM-PP1 | ÜB44007 | 63 |
